# Topology-Encoded Polarity in Oppositely Charged Binary IDPP Condensates: Multiphase Organization from Non-Coacervating Partners as a Minimal Model of Complex Coacervation

**DOI:** 10.64898/2025.12.05.692546

**Authors:** Julio Fernández-Fernández, Vicente Domínguez-Arca, Raúl Escribano, Sergio Ferrero, Sergio Acosta, Jose Carlos Rodríguez-Cabello

## Abstract

Synthetic condensates provide a way to engineer compartmentalized microenvironments that mimic the properties and functions of natural ones, yet the principles that govern their phase behavior and internal organization remain incompletely defined. Introducing charged residues into intrinsically disordered protein polymers (IDPPs) with LCST phase behavior typically suppresses phase separation under physiological conditions. Here we show that pairing two such oppositely charged IDPPs restores and programs LCST-driven liquid-liquid phase separation (LLPS), enabling a minimalist two-component platform for constructing synthetic condensates whose formation, size, and internal organization are encoded directly in sequence. LLPS emerges from an asymmetric, entropy-driven interplay between hydrophobic collapse, solvent reorganization, and salt-bridge topology. The balance between inter- and intrachain ionic pairing leads to distinct dense-phase microenvironments with tunable residual charge and micropolarity, thereby controlling condensate formation, and miscibility and the emergence of single-phase or multiphase protein condensates. The condensate interior further alters the ionization thermodynamics of charged residues shifting their apparent pKa and enabling tunable pH responses. Systems dominated by interchain salt bridges form low-polarity condensates that mix uniformly with hydrophobic partners, whereas molecular architectures favoring intrachain pairing retain residual charge and, in the presence of hydrophobic partners, undergo spontaneous internal demixing into multiphase assemblies. These findings establish a mechanistic, sequence-level framework for encoding phase behavior, micropolarity, and mesoscale organization in synthetic condensates, and demonstrate how minimalistic LCST-IDPP pairs can be engineered to create programmable microenvironments, opening avenues toward engineered condensates with higher-order organization and adaptive capabilities.

## 1. Introduction

Supramolecular assemblies generated from short peptides and protein-engineered polymers have become a powerful and chemically minimalist platform for bottom-up biofabrication of functional biomaterials and bioactive interphases.[1,2] In many of these systems, structural motifs such as β-sheets, coiled coils, or α-helices provide well-defined secondary and even tertiary organization, enabling the formation of β-sheet fibrils, [3] nanorribons,[4,5] protein cages [6], giant vesicles [7], liquid droplets or shear-thinning hydrogels.[8] These “ordered” supramolecular assemblies dominate the field and are valued for their predictability and robust mechanical properties, although their stability often comes at the cost of irreversible aggregation, as seen in amyloid fibrils.

In contrast, intrinsically disordered proteins (IDPs) and their synthetic analogues, intrinsically disordered protein polymers (IDPPs), lack persistent secondary structure. Their assemblies arise instead from dynamic, multivalent interactions that resemble biomolecular condensates in cells.[9] This complementary “disordered” design strategy is emerging as a versatile alternative, offering reversible phase behavior, tunable responsiveness, and biocompatibility, thereby expanding the design space of supramolecular biofabrication.[10,11] Within this broader context, two-component and multicomponent systems provide an additional hierarchical level: by combining ordered and/or disordered motifs, orthogonal functionalities can be encoded in each assembling partner while cooperative interactions drive the emergence of properties that neither component displays individually.[12] Such cooperative design and organization principles are now being harnessed in peptide- and protein-based biomaterials for catalysis, sensing, soft-robotics, vaccine delivery, and regenerative medicine. [13–16]

This dichotomy between ordered and disordered assemblies is directly mirrored in biology, where IDPs exemplify how dynamic molecular organization can be exploited in living systems.[17] Lacking a fixed secondary structure, they adapt rapidly to fluctuating environments, allowing cells to regulate biochemical processes spatiotemporally. A central mechanism underlying this regulation is liquid–liquid phase separation (LLPS), through which IDPs condense into biomolecular condensates or membraneless compartments. These biomolecular condensates transiently concentrate small molecules and macromolecules, thus controlling buffer fluctuations, [18,19] generating electrochemical potentials, or coordinate signaling pathways,[20] illustrating how structural disorder serves as a functional advantage rather than a deficiency.

Building upon this biological paradigm, IDPPs have been developed as synthetic analogues that not only recapitulate but also extend the LLPS phenomena observed in native IDPs.[10,21] Their genetic programmability and sequence tunability enable the construction of supramolecular systems with finely tailored responsiveness. The driving forces governing IDPP condensation mirror those of their natural counterparts,[22] and can be broadly divided into two regimes.[10] Resilin-like sequences (e.g., GRGDSPYS), enriched in aromatics and charged residues, rely on π–π, cation–π, electrostatic, and hydrophobic interactions, often giving rise to UCST-type behavior. By contrast, elastin-like repeats such as VPGXG primarily undergo hydrophobic collapse, leading to LCST-mediated condensation. These two regimes thus represent the enthalpy- and entropy-dominated extremes of protein phase separation, respectively. Strategically combining electrostatic complementarity with LCST-driven desolvation provides a powerful strategy to control not only phase separation thresholds but also the internal organization of IDPP condensates.

Synthetic condensates built from IDPPs are increasingly being explored as functional modules that mimic cellular functions. Such assemblies can create catalytic microenvironments, regulate metabolic fluxes, or serve as programmable compartments for sequestration and release of biomolecules. By concentrating enzymes with their substrates, they can enhance catalytic efficiency in a manner reminiscent of stress-induced condensates.[23] Incorporating nucleic acid-binding domains into IDPPs enables the selective sequestration of plasmids and mRNA, thereby modulating gene expression or protecting from degradation in vivo.[24,25] Moreover, tuning the amino-acid composition or “molecular grammar” of IDPPs allows precise tuning of condensate properties,[26,27] including phase behavior,[28] interfacial electrochemical potential,[19] viscoelasticity,[29] and micropolarity,[30] a recently uncovered determinant of multilayered condensate organization.

Beyond the intrinsic sequence grammar that governs single-component condensates, biological LLPS almost always arises from both homotypic and heterotypic interactions. [31,32] In living cells, condensate stability and composition are not fixed properties but emerge from the thermodynamics of multicomponent mixtures, where heterotypic associations between distinct biomolecules typically provide the cohesive forces that stabilize condensates, whereas homotypic associations modulate their selectivity and internal organization.[31] Translating this principle to synthetic systems introduces an additional layer of design complexity. The balance between homotypic associations within each protein and heterotypic interaction between distinct partners can modulate condensate miscibility, internal organization, and responsiveness, as shown in both short peptide and IDPP-based condensates. [30,33,34] Understanding how this competition shapes phase behavior is therefore critical to rationally design bicomponent IDPP condensates with programmable architectures and dynamic physicochemical properties.

Despite these advances, no synthetic condensates have yet been constructed exclusively from bicomponent IDPP systems. Establishing design rules for such assemblies is both a mechanistic necessity and a transformative opportunity. In this study, we elucidate the critical role of electrostatic complementarity and molecular architecture in the co-assembly of oppositely charged IDPPs. Through a combination of calorimetries, confocal microscopy, turbidity assays, circular dichroism, nuclear magnetic resonance (NMR), and atomistic simulations, we dissect the energetic variables that govern condensate formation, providing a conceptual and quantitative framework for the rational design of functional protein condensates from two-component systems. We show that charge distribution and sequence architecture dictate not only phase behavior but also the balance between intra- and intermolecular salt bridges, with direct consequences for condensate properties. Furthermore, we demonstrate that condensate local environment shifts protonation equilibria and enables subcompartmentalization when combined with hydrophobic IDPPs. Together, these findings position bicomponent IDPP condensates as both reductionist models to probe the molecular grammar of natural IDPs and as programmable synthetic organelles whose phase behavior and internal chemistry can be directly encoded in sequence.

## 2. Results and discussion

### 2.1. Co-assembly in solution

To establish a minimalist platform to probe co-assembly of oppositely charged IDPPs, we engineered a library of LCST-IDPPs based on the canonical elastin-like pentapeptide motif, VPGVG (**Figure 1a, Tables S1-S2, Figures S1-S2)**. In these constructs, one in every five pentapeptides was substituted, at the X position, with either glutamic acid (Glu, E) or lysine (Lys, K), generating two oppositely charged 50-mer (50 pentamers) recombinant polypeptides, designated V_2_EV_2_ and V_2_KV_2_, respectively. This design preserved the hydrophobic backbone responsible for LCST behavior while introducing a controlled density of ionizable residues. The molecular architecture of both IDPPs is essentially identical being one the exact specular image, in terms of charge sign, of the other. The resulting IDPPs were thus expected to display competing electrostatic interactions: repulsive homotypic contacts within chains carrying like charges (V_2_KV_2_ + V_2_KV_2_ or V_2_EV_2_ + V_2_EV_2_), and efficient attractive heterotypic interactions between oppositely charged partners (V_2_KV_2_ + V_2_EV_2_). We hypothesized that the balance between these opposing forces would govern the onset and reversibility of the LLPS. To further dissect the role of sequence architecture, we also constructed an IDPP in which both charged polypeptides were covalently linked in a diblock copolymer (V_2_KV_2_-V_2_EV_2_). Additionally, two non-charged analogues, (VPGVG)_50_ (V) and (VPGVG)_100_ (V-V) were also produced.

**Figure 1.**
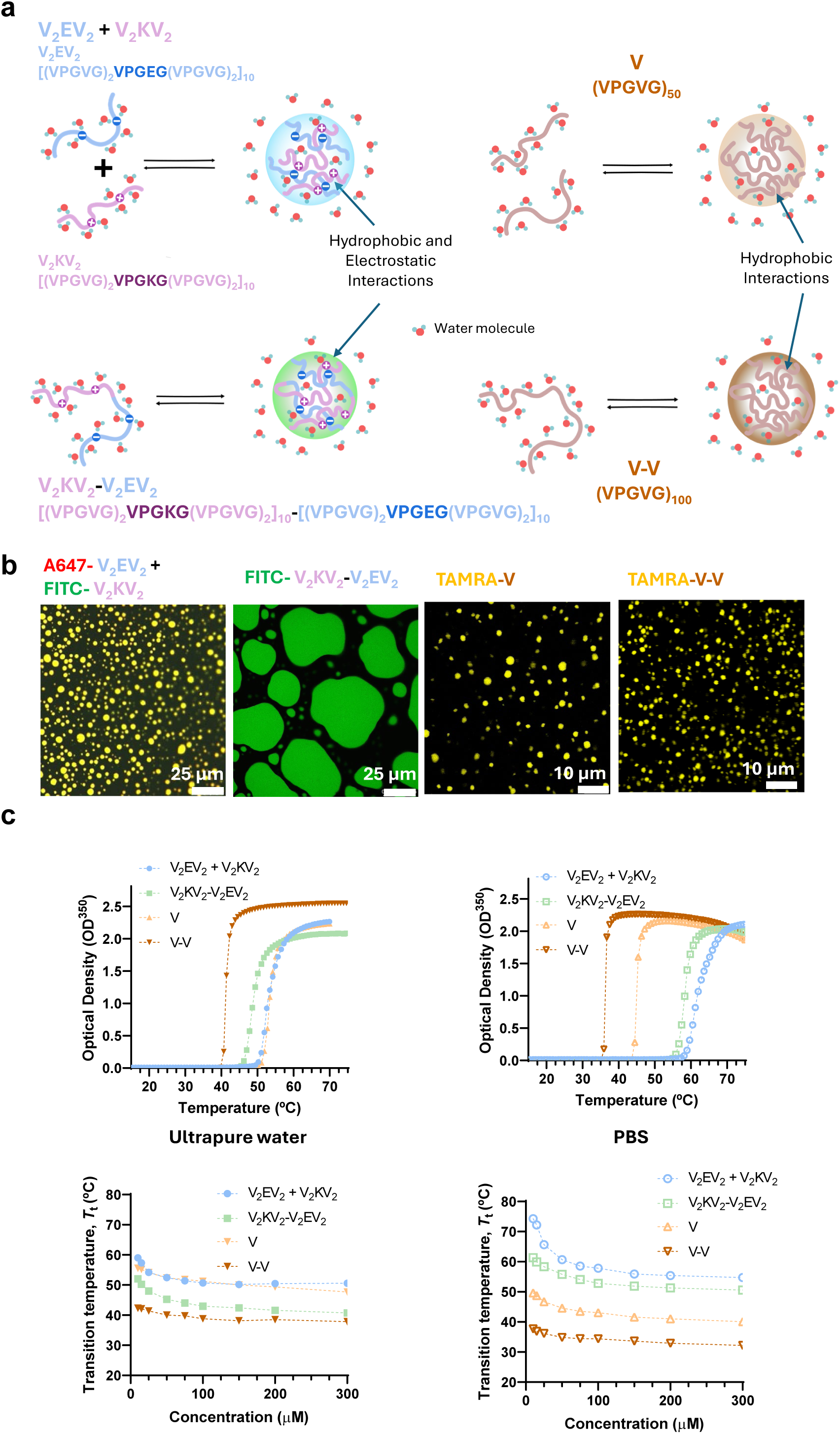
Sequence architecture dictates LLPS and condensate morphology. a) Schematic representation of five IDPPs and their LLPS in aqueous solution: the equimolar mixture (V_2_EV_2_ + V_2_KV_2_) of two oppositely charged IDPPs (i.e., the cationic V_2_KV_2_ [(VPGXG)_50_, X = Val/Lys 4:1] and the anionic V_2_EV_2_ [(VPGXG)_50_, X = Val/Glu 4:1]) diblock copolymer (V_2_KV_2_-V_2_EV_2_), and two neutral variants, (VPGVG)_50_ (V) and (VPGVG)_100_ (V-V). Charge complementarity in V_2_KV_2_/V_2_EV_2_ systems enables both electrostatic and hydrophobic interactions during condensation, whereas VPGVG-based homopolymers undergo phase separation solely through hydrophobic collapse. b) Confocal micrographs of the condensates formed with fluorescent labelled IDPPs. All IDPPs generate spherical condensates except V_2_KV_2_-V_2_EV_2_, which forms larger, irregular, patch-like domains indicative of altered phase separation dynamics. Scale bars: 25 μm (V_2_EV_2_ + V_2_KV_2_ and V_2_KV_2_-V_2_EV_2_, left), 10 μm (V and V-V, right) c) UV/Vis turbidimetry profiles and d) phase diagrams in ultrapure water and PBS reveal the impact of ionic strength in modulating LLPS. In charged systems (V_2_EV_2_ + V_2_KV_2_ and V_2_KV_2_-V_2_EV_2_), electrostatic screening increases the cloud point temperature (*T*_t_), consistent with reduced effective attraction upon charge neutralization. In contrast, in hydrophobic IDPPs (V and V-V), salt lowers *T*_t_, indicating enhanced hydrophobic dehydration and chain collapse. IDPP concentrations in (c) were normalized to the number of pentapeptide repeats: 25 μM for V-V and V_2_KV_2_-V_2_EV_2_, and 50 μM for V_2_EV_2_ + V_2_KV_2_ and V.

Individually, and as expected, V_2_EV_2_ and V_2_KV_2_ showed no measurable cloud point in water at pH 7 (**Figure S3**), consistent with strong electrostatic repulsion and the enhanced polarity conferred by ionizable residues, which broadens their solubility window across temperatures.[35] In contrast, the equimolar V_2_EV_2_ + V_2_KV_2_ mixture exhibited a clear reversible phase separation (**Figure 1c**), producing protein-rich microdroplets (**Figure 1b**) above a well-defined cloud point transition temperature (*T*_t_), nearly identical to that of the uncharged analogue V (**Figure 1c, 1d**). This similarity in thermal behavior in water, together with the droplet morphology, suggests that charge neutralization within the dense phase in the binary system is nearly complete, such that LLPS is primarily driven by hydrophobic collapse.

The diblock IDPP, V_2_KV_2_-V_2_EV_2_, also underwent a well-defined LCST-type LLPS, yet with altered demixing dynamics compared with the two-component system. Immediately above *T*_t_, spherical droplets nucleated in solution but coalesced within minutes into irregular patch-like condensates (**Figure 1b**). V_2_KV_2_-V_2_EV_2_ underwent LLPS at a lower *T*_t_ than the V_2_EV_2_/V_2_KV_2_ mixture (**Figure 1c,d**). This behavior would be, initially, consistent with the general dependence of IDPPs on chain length, whereby increasing molecular weight lowers *T*_t_ due to cooperative dehydration. [36] However, the reduction was less pronounced than in the non-charged analogue (V-V), despite both polymers sharing the same total number of pentapeptides. This attenuation suggests that intramolecular charge compensation in V_2_KV_2_-V_2_EV_2_ partially counterbalances the chain-length effect, thereby constraining the extent to which molecular weight alone can depress the *T*_t_.

At physiological ionic strength, i.e., in PBS, the phase behavior of the IDPP library varied depending on the presence or absence of charges. In charged systems (V_2_EV_2_ + V_2_KV_2_ and V_2_KV_2_-V_2_EV_2_), *T*_t_ increased compared to that found in ultrapure water, consistent with electrostatic screening that weakens attractive interactions. In contrast, the uncharged IDPPs (V and V-V), displayed the opposite trend, with salt lowering *T*_t_, due to the well-known salting-out effect that promotes hydrophobic dehydration and chain collapse.[37] Notably, although V-V and V_2_KV_2_-V_2_EV_2_ contain the same number of pentapeptide repeats, their chain-length effects on *T*_t_ were not equivalent. In both cases, increasing molecular weight lowered the *T*_t_ relative to their shorter analogues, consistent with the cooperative dehydration typical of LCST IDPPs, whereby longer chains promote more extensive desolvation and aggregation.[38] However, this reduction was markedly attenuated in the charged diblock, suggesting that the electrostatic component partially disrupts the cooperative effect. One possible explanation is that charge neutralization within V_2_KV_2_-V_2_EV_2_ is incomplete, leaving a residual net charge that enhances hydration and increases the *T*_t_. Alternatively, intramolecular charge pairing within the covalently linked blocks alters the spatial distribution of exposed polar and charged residues, making it distinct from the intermolecular interactions that occur in the V_2_EV_2_ + V_2_KV_2_ mixture.

### 2.2. Two energy landscapes revealed by ionic strength and molecular architecture

To dissect the energetic basis of V_2_EV_2_/V_2_KV_2_ co-assembly, we analyzed the thermal transitions of mixtures at varying stoichiometries by differential scanning calorimetry (DSC) (**Figure 2a, c** and **S4a**). The observed endotherms report on the energetic cost of disrupting protein-water hydrogen bonds and releasing hydration layers and counterions, compensated by favorable protein-protein interactions in the protein-rich phase. Accordingly, higher transition enthalpies (ΔH) reflect more extensive molecular reorganization during LLPS.

**Figure 2.**
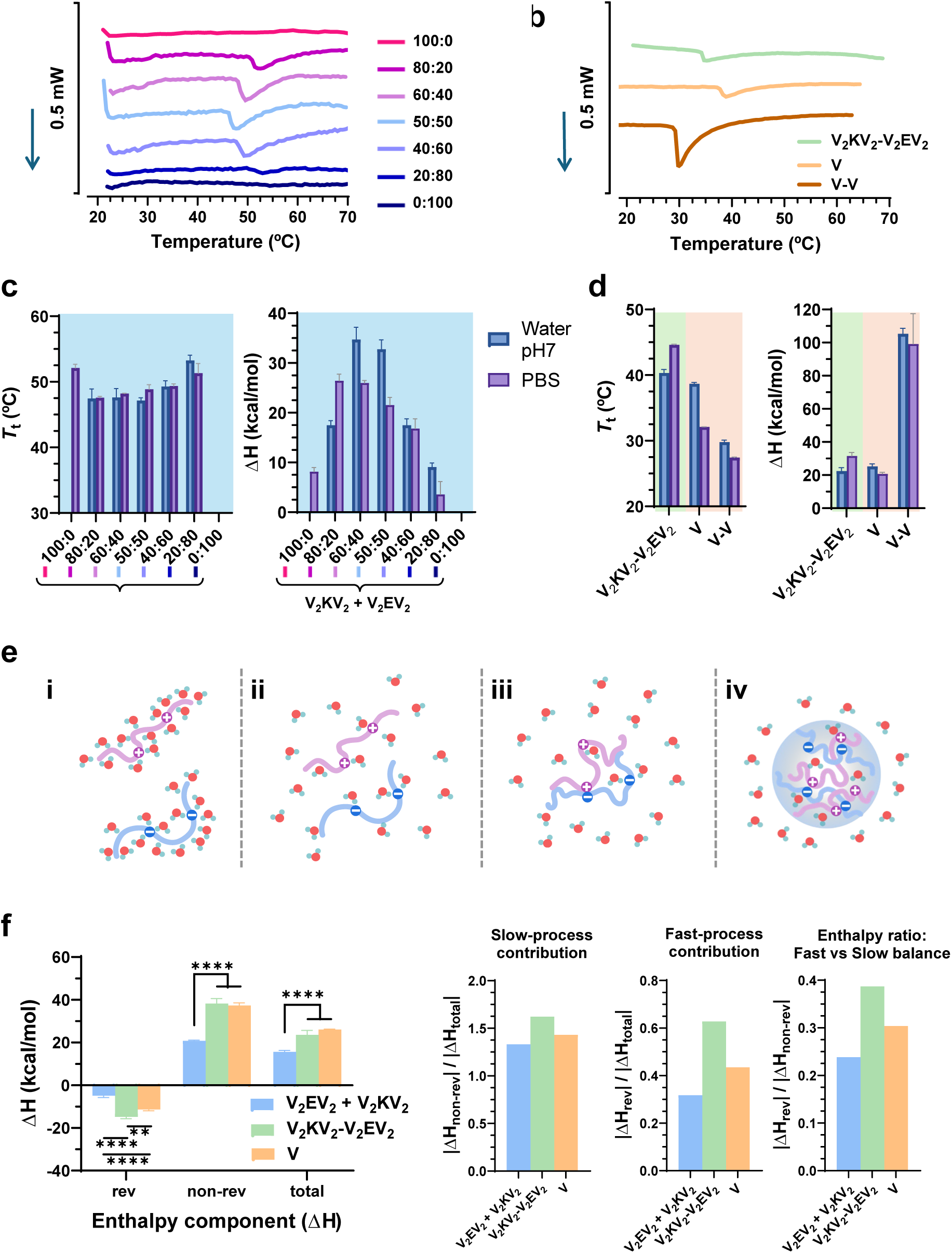
Thermodynamic evaluation reveals the influence of charge stoichiometry, protein architecture and hydrophobicity on the LLPS. (a, b) Conventional DSC thermograms of (a) mixtures of V_2_KV_2_ and V_2_EV_2_ at different molar ratios (100:0 to 0:100), and (b) of the diblock copolymer V_2_KV_2_- V_2_EV_2_ and the hydrophobic IDPPs V and V-V reveal distinct endothermic profiles depending on the molecular architecture. (c, d) Transition temperatures (*T*_t_) and total enthalpy changes (ΔH_tot_) extracted from conventional DSC data plotted for (c) mixtures of V_2_KV_2_ + V_2_EV_2_ at varying charge ratios and (d) the V_2_KV_2_-V_2_EV_2_, V, V-V systems. The LLPS in the binary V_2_KV_2_ + V_2_EV_2_ system shows a clear asymmetric dependence on composition, where maximum exothermic enthalpy release occurs at 60:40 V_2_KV_2_:V_2_EV_2_. (e) Schematic representation of the condensation mechanism. (i) Below the *T*_t_, the IDPP chains are hydrophobically hydrated forming clathrate-like structures. (ii) Upon heating above *T*_t_ (LCST), IDPP chains undergo cooperative desolvation of structured water, followed by (iii) chain collapse and intra-and intermolecular interactions, culminating in (iv) condensate formation via LLPS. (f) Deconvolution of enthalpy components via TM-DSC. Reversing component (ΔH_rev_) corresponds to fast, cooperative and predominantly endothermic processes, specifically the formation of hydrophobic and electrostatic interactions that drive phase separation (i.e., protein-protein interactions). The non-reversing component (ΔH_non-rev_) arises from slower processes, such as the diffusion and release of structured water molecules (i.e., clathrate-like) which are physically reversible but occurs on longer timescales. (g) Enthalpic ratios derived from TM-DSC deconvolution highlight the relative weight of fast and slow processes in LLPS. Left: ΔH_non-rev_/ΔH_total_ represents the contribution of slow processes. Center: ΔH_rev_/ΔH_total_ reports the contribution of fast-processes, i.e., the fraction of the transition enthalpy compensated by rapid reversing events. Right: ΔH_rev_/ΔH_non-rev_ indicates the ratio between both components.

In ultrapure water (pH 7), neither V_2_EV_2_ nor V_2_KV_2_ alone underwent LLPS, whereas all the mixtures of V_2_EV_2_ and V_2_KV_2_ underwent endothermic LLPS in the range of 9-35 kcal mol^-1^ depending on the stoichiometry. The largest ΔH value was observed not at the equimolar ratio but at 60:40 V_2_KV_2_: V_2_EV_2_ (ΔH_60:40_ = 34.7 ± 2.5 kcal·mol^-1^), slightly higher than the equimolar mixture (ΔH_50:50_ = 32.8 ± 1.9 kcal mol^-1^). This asymmetric maximum suggests that Lys and Glu do not contribute equally to electrostatic neutralization. The localized negative charge of Glu carboxylate (-COO^-^) might interact with multiple Lys side chains, consistent with systems based on polyelectrolyte complexes and charge-complementary peptides.[39,40]

In PBS, the enthalpic profile of V_2_EV_2_/V_2_KV_2_ mixtures varied in comparison to water, highlighting the influence of ionic strength on the balance between charge compensation and dehydration (**Figure 2c and S4a**). Unbalanced V_2_KV_2_-rich mixtures (80:20 and 60:40) showed the highest ΔH values (e.g., ΔH_80:20_ = 26.4 ± 1.4 kcal·mol^-1^ and ΔH_60:40_ = 26.0 ± 0.5 kcal·mol^-1^), whereas V_2_EV_2_-rich systems exhibited strongly reduced values (ΔH_40:60_ = 16.8 ± 2.0; ΔH_20:80_ = 3.6 ± 2.6 kcal·mol^-1^). Remarkably, pure V_2_KV_2_ (100:0), which remained fully soluble in water, underwent LLPS in PBS (ΔH_100:0_ = 8.15 ± 0.83 kcal·mol^-1^). This behavior cannot be explained solely to electrostatic screening of repulsive Lys-Lys interactions but likely reflects the combined influence of charge screening and salting-out. The ionic strength reduces the hydration capacity of Lys ammonium groups (-NH_3_^+^), which solvation free energy is lower than Glu carboxylates,[41] thereby unmasking a latent hydrophobic collapse. In contrast, V_2_EV_2_ remained soluble in PBS, even at high concentrations (5% w/v), consistent with robust hydration of carboxylate groups.

Molecular architecture modulated these trends. In water, V_2_KV_2_-V_2_EV_2_ underwent LLPS at lower *T*_t_ and ΔH 40.3 °C; 22.4 ± 2.1 kcal·mol^-1^) than V_2_EV_2_ + V2KV2 (47.1 °C; 32.8 ± 1.9 kcal·mol^-1^), consistent with intrachain charge pairing that may partially frustrate cooperative dehydration. In PBS, however, the behavior inverted, and V_2_KV_2_-V_2_EV_2_ exhibited a significantly higher ΔH (31.6 ± 2.1 kcal·mol^-1^) and a stronger shift in *T*_t_ (44.6 °C) compared to the equimolar mixture. This inversion suggests that under physiological ionic strength, charge screening may impair intermolecular electrostatic attraction in the two-component system, but intramolecular salt bridges within V_2_KV_2_-V_2_EV_2_ system are less affected by bulk counterions and the salting-out effect is enhanced at high concentrations.

Non-charged IDPPs (V and V-V) followed canonical LCST behavior. V exhibited a *T*_t_ significantly lower than the V_2_EV_2_/V_2_KV_2_ mixture in water, with an ΔH of 25.3 ± 1.5 kcal·mol^-1^. Doubling the chain length to V-V further decreased *T*_t_ and produced a non-linear increase in enthalpy to 105.4 ± 3.2 kcal·mol^-1^, consistent with previous studies on VPGVG polymers.[42] In PBS, both underwent salting-out, lowering their *T*_t_ relative to water. Their enthalpies again differed with chain length, consistent with stronger cooperative dehydration at higher molecular weights.

#### Thermodynamic dissection of LCST-type phase separation

Phase separation in LCST-type IDPP condensates is governed by two thermodynamically coupled yet kinetically distinct processes: a slow, cooperative dehydration accompanied by counterion release and a faster molecular reorganization within the emerging dense phase (schematically represented in **Figure 2e**).[42] This hierarchical coupling between solvent reorganization and chain rearrangements is consistent with the general physical principles described for intracellular phase transitions, where enthalpy-driven association and solvent reorganization operate on different timescales.[43] The resulting energetic partitioning defines two complementary contributions to the overall LLPS in our system: (i) a cooperative, enthalpically costly step associated with the expulsion of structured water from clathrate-like hydration shells during hydrophobic desolvation and electrostatic neutralization, and (ii) a rapid, compensatory step that provides enthalpic and entropic recovery through local chain reorganization and salt-bridge formation.

To experimentally resolve these coupled processes, we employed temperature-modulated DSC (TM-DSC) to decompose the total transition enthalpy (ΔH_tot_) into its non-reversing (slow) and reversing (fast) components (**Figure 2f, g**). The non-reversing term (ΔH_non-rev_) corresponds to the cooperative dehydration and counterion release, whereas the reversing term (ΔH_rev_) captures rapid, reversible rearrangements that transiently offset this enthalpic cost.

Across all systems, ΔH_non-rev_ dominated the condensate formation, yielding positive ΔH_tot_ values consistent with conventional DSC. Quantitatively, the non-charged IDPP, V, showed a large slow component (ΔH_non-rev_ = 37.4 ± 1.2 kcal·mol^-1^) partially compensated by a modest reversing (fast) exotherm (ΔH_rev_ = -11.4 ± 0.6 kcal·mol^-1^), resulting in a ΔH_total_ = 26.1 ± 0.2 kcal·mol^-1^. V_2_KV_2_-V_2_EV_2_ displayed similar slow term (ΔH_non-rev_ = 38.2 ± 2.4 kcal·mol^-1^), but a stronger fast exotherm (ΔH_rev_ = - 14.8 ± 0.8 kcal·mol^-1^), reducing ΔH_total_ to 23.6 ± 2.1 kcal·mol^-1^. In contrast, the equimolar V_2_EV_2_ + V_2_KV_2_ mixture showed the smallest enthalpies, with ΔH_non-rev_ = 20.7 ± 0.4 kcal·mol^-1^ and ΔH_rev_ = -4.9 ± 0.8 kcal·mol^-1^ (ΔH_total_ = 15.6 ± 0.7 kcal·mol^-1^).

These results reveal how molecular architecture and charge distribution modulate the relative contributions of slow and fast enthalpic components. Uncharged IDPPs (V) undergo LLPS primarily through slow, cooperative dehydration whereas the covalently-bonded V_2_KV_2_-V_2_EV_2_ exhibit a larger reversing (fast) contribution, consistent with intrachain electrostatic neutralization that might promote rapid reorganization prior to macroscopic phase separation. In this architecture, oppositely charged domains within the same chain can establish transient intramolecular salt bridges, effectively reducing repulsive forces and facilitating the subsequent hydrophobic coacervation that drives LCST-type condensation.

Conversely, in the V_2_EV_2_ + V_2_KV_2_ mixture, charge neutralization can only occur through intermolecular interactions between independently diffusing chains. These encounters are stochastic and spatially diffuse and may lead to a less efficient compensation of electrostatic repulsion. As a result, dehydration proceeds in a less cooperative and energetically weaker manner, reflected in the lower ΔH_non-rev_ and ΔH_rev_ values.

Analysis of enthalpy ratios further emphasizes these differences (**Figure 2g**). In all systems, slow cooperative processes dominate (ΔH_non-rev_ / ΔH_total_ > 1), confirming that slow processes, associated mainly with dehydration and counterion release, dominate in the LLPS. However, the degree of fast contribution varied markedly with ratios of ∼43% for V, ∼63% for V_2_KV_2_- V_2_EV_2_, and minimal for V_2_EV_2_ + V_2_KV_2_ (∼32%). This trend supports that stoichiometry and molecular architecture not only determine the total enthalpy of LLPS but also dictates its kinetic partitioning. The enhanced fast contribution in V_2_KV_2_-V_2_EV_2_ reflects the effect of intrachain electrostatic reorganization and salt-bridge formation, which provide rapid enthalpic compensation absent in purely hydrophobic systems. By contrast, the attenuated fast contribution in the free V_2_EV_2_ + V_2_KV_2_ mixture suggests that when salt bridge formation only occurs interchain, it might lead to a less organized manner. This would explain why ΔH_total_ is lowest in the mixture despite the presence of both charged residues, while V and V_2_KV_2_-V_2_EV_2_ exhibit stronger cooperative reorganization that translates into higher ΔH_total_. Consistently, these enthalpic differences align with the distinct secondary structures observed across condensates (**Figures S5** and **S6**).

Control experiments performed at strongly acid and basic conditions further clarified this picture (**Figure S7**). Below and above their respective pK_a_ values (i.e., pK_aE_ and pK_aK_), both V_2_EV_2_ and V_2_KV_2_ lost electrostatic charges and behaved as non-charged IDPPs with clear endotherms. V_2_KV_2_ at pH 12 displayed ΔH_total_ = 27.7 ± 0.3 kcal·mol⁻¹ (dominated by ΔH_non-rev_ = 33.8 ± 1.1). V_2_EV_2_ at pH 3 showed an even larger enthalpic value (ΔH_total_ = 42.2 ± 0.1, ΔH_non-rev_ = 49.6 ± 1.6), consistent with stronger cooperative dehydration of protonated carboxylic groups compared to lysine amine groups. These results confirm that the asymmetric contribution of charged residues explains the distinct behavior of V_2_EV_2_/V_2_KV_2_ mixtures.

### 2.3. Two-component IDPP system undergo asymmetric, entropy-governed condensation

To connect the calorimetric energy partition with molecular interactions, we turned to isothermal titration calorimetry (ITC) and atomistic molecular dynamics (MD). ITC quantifies binding stoichiometry and cooperativity within the condensed state, while MD visualizes chain compaction, salt-bridge formation, and counterion release across *T*_t_. Together, these measurements test whether the asymmetric enthalpy maximum and architecture-dependent ΔH_rev_ contributions observed by DSC originate from unequal charge neutralization and collective solvent reorganization at the molecular scale.

To determine the molecular affinity and stoichiometry driving V_2_EV_2_ + V_2_KV_2_ co-assembly, we performed ITC experiments in both directions, direct (V_2_EV_2_→V_2_KV_2_) and reverse (V_2_KV_2_→V_2_EV_2_), at temperatures below and above the *T*_t_ of the mixture (**Figure 3**). Above the *T*_t_, both titrations displayed an abrupt decay of the calorimetric signal with molar ratio, indicative of highly cooperative, multivalent protein-protein interactions within the two-component condensate. Fitting of the binding isotherms (details in **Supporting information**) enabled extraction of the stoichiometry *n*, binding constant *K*, and enthalpy ΔH for both titration directions. For V_2_EV_2_→V_2_KV_2_, the best-fit values were Δ*H* = 8.8 × 10^5^ J mol*^−^*^1^ (210.33 kcal mol*^−^*^1^), *K* = 4.1 × 10^9^ M*^−^*^1^, and *n* = 0.55. For V_2_KV_2_→V_2_EV_2_, the fitted parameters were Δ*H* = 5.4 × 10^5^ J/mol (129.06 kcal mol^-1^), *K* = 9.1 × 10^9^ M*^−^*^1^, and *n* = 1.36. In other words, although both titrations exhibit strong binding affinities and similar enthalpic signatures, the effective stoichiometries diverge markedly. Condensation requires approximately twice as much V_2_KV_2_ as V_2_EV_2_ to reach saturation, in line with the enthalpy maximum shifted toward V_2_KV_2_-rich mixtures observed by DSC.

**Figure 3.**
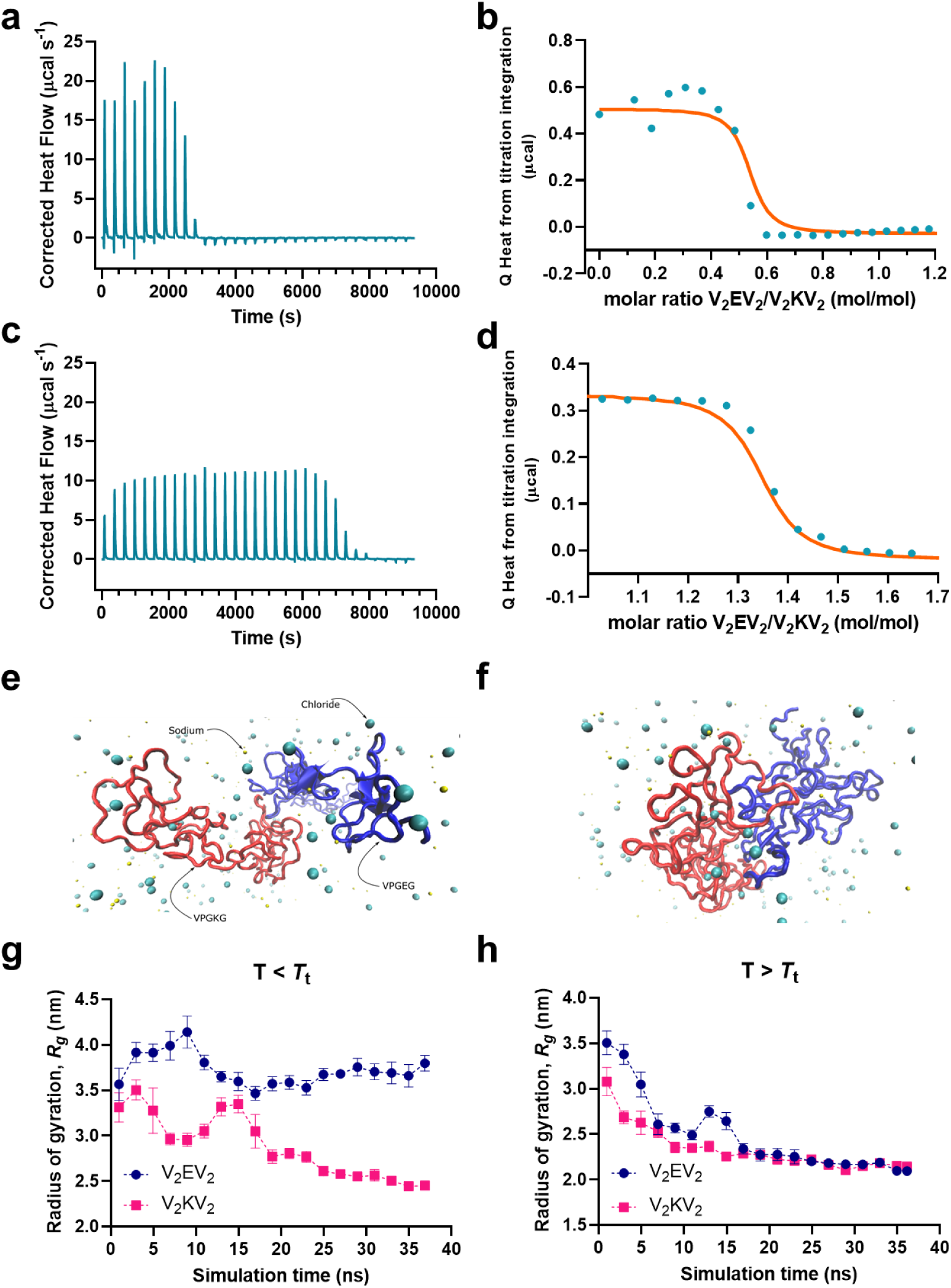
Thermoresponsive LLPS between V_2_EV_2_ and V_2_KV_2_ as revealed by ITC and molecular simulations. (a) Raw calorimetric data with baseline correction for V_2_EV_2_→V_2_KV_2_ titration above the *T*_t_ (T = 60°C). (b) Corresponding binding isotherm and nonlinear fit. (c) Raw calorimetric data for V_2_KV_2_→V_2_EV_2_ titration above the *T*_t_ (T = 60°C). (d) Binding isotherm and fit. (e) Final snapshot of the V_2_EV_2_-V_2_KV_2_ mixture below the *T*_t_ (T = 20°C) showing spatial separation and solvation shells (Na^+^, Cl^-^ identified). (f) Final snapshot at 60◦C showing compacted, interacting IDPPs and ion redistribution. (g) Radius of gyration (*R_g_*) for V_2_EV_2_ and V_2_KV_2_ below the *T*_t_ (T = 20°C) indicating limited structural collapse. (h) *R_g_* above the *T*_t_ (T = 60°C), showing ∼4 °A decrease, consistent with thermal condensation.

**Figure 4.**
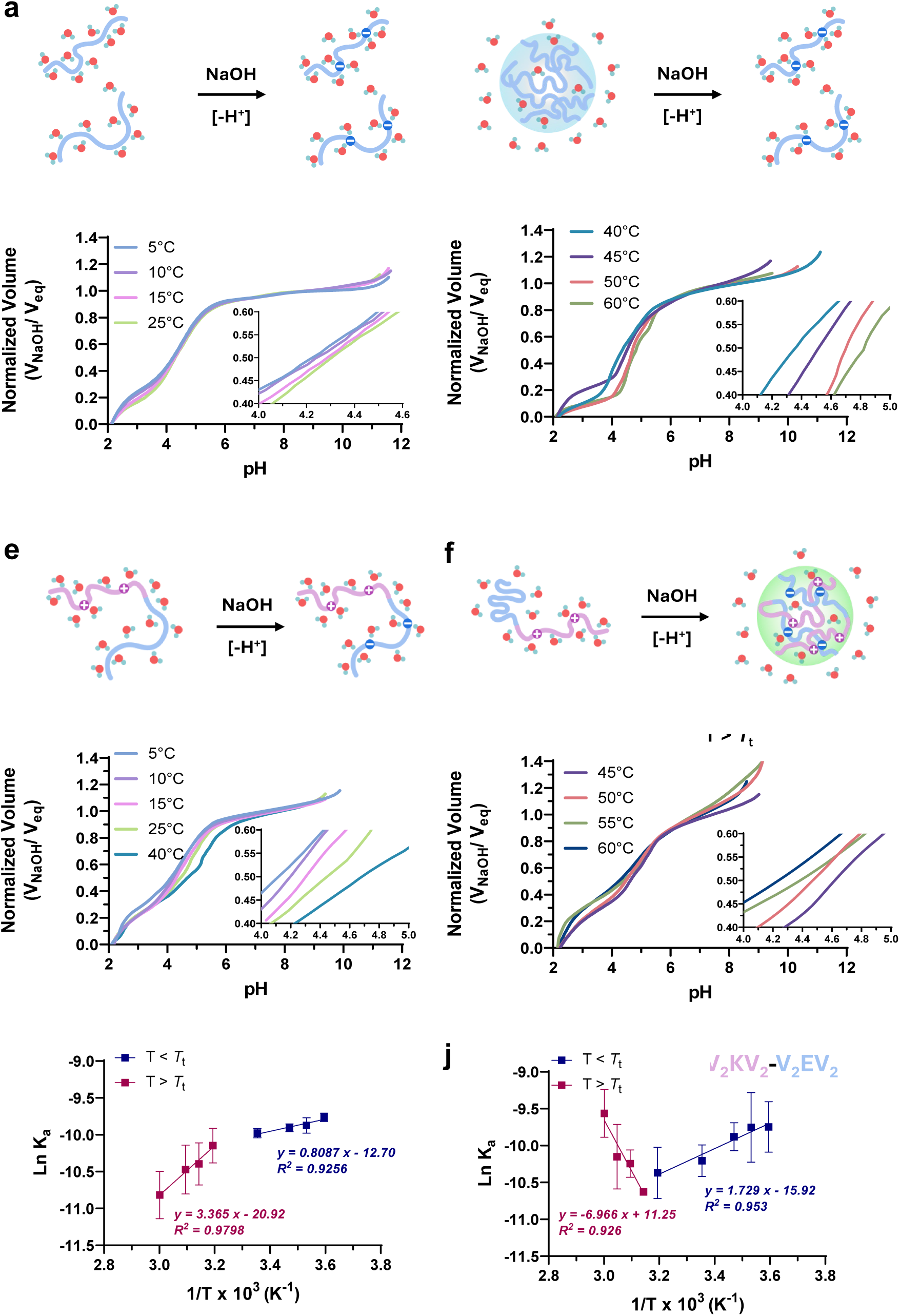
Thermodynamic characterization of the LLPS of V_2_EV_2_ and V_2_KV_2_-V_2_EV_2_ by acid–base titration above and below the *T*_t_. Schematic representations of the phase behavior and protonation state of (a, b) V_2_EV_2_ or (e, f) V_2_KV_2_-V_2_EV_2_ titrated below (a, e) or (b, f)) above their *T*_t_. V_2_EV_2_ undergoes condensation only when it is protonated (uncharged, below pK_a_) and heated above LCST, whereas V_2_KV_2_-V_2_EV_2_ remains soluble at high temperature until deprotonation triggers LLPS. (c, d, g, h) Representative titration curves for V_2_EV_2_ (c, d) and V_2_KV_2_-V_2_EV_2_ (g, h) monitored across multiple temperatures below (c, g) and above (d, h) *T*_t_. Insets zoom into the pK_aE_ region (0.5 V_eq_), highlighting the shift in protonation equilibria upon phase separation. (i–j) Van’t Hoff plots constructed from titration equilibrium constants at different temperatures for VE (i) and V_2_KV_2_-V_2_EV_2_ (j), showing linear relationships that allow estimation of apparent ΔH° and ΔS° values. V_2_KV_2_-V_2_EV_2_ exhibits an inverse enthalpic trend compared to V_2_EV_2_ above the LCST, consistent with a condensation mechanism driven by charge neutralization and hydrophobic collapse and modulated by the amphiphilic diblock architecture.

Interestingly, the magnitude of these enthalpic changes differs markedly from those reported for natural IDP-IDP interactions and complex protein coacervates. The coupling enthalpies of IDP complexes typically fall within the range of 0.1–10 kcal·mol^-1^, [44,45] reflecting local rearrangements within a single binding interface and limited solvent reorganization.[39] Oppositely charged polypeptides exhibit even smaller enthalpic signatures,[46] on the order of 1–3 kJ·mol⁻¹ (0.2–0.7 kcal·mol⁻¹), as the process is largely entropy-driven by counterion and water release. In contrast, the enthalpies measured in our system —on the order of 10^5^ J·mol^-1^ (tens to hundreds of kcal·mol^-1^ per polymer)—are two to three orders of magnitude higher than those of IDPs, and three to four orders of magnitude higher than those of classical polyelectrolyte complex coacervation. These unusually high values do not represent single binding events between residues; rather, they capture the cumulative, cooperative reorganization of hundreds of weak, transient interactions and extensive solvent restructuring occurring during LCST-driven condensation.

To visualize this collective molecular reorganization and connect thermodynamics with nanoscale structure, atomistic MD simulations of equimolar V_2_EV_2_ + V_2_KV_2_ mixtures were performed at temperatures just below and above the *T*_t_. Below the *T*_t_, both IDPPs remained solvated and spatially separated, with intact ion clouds of Na^+^ and Cl*^−^* surrounding the chains, especially around the charged side chains. In contrast, above the *T*_t_, the frame showed tight interchain association and partial expulsion of counterions from the condensed chains. This reorganization coincided with a ∼4 Å decrease in the radius of gyration (*R_g_*) for both chains (**Figure 3g, h**), indicating thermally induced compaction consistent with the ITC results. The concomitant decrease in solvent-accessible surface area (SASA) further confirmed the loss of solvation shell and the burial of hydrophilic and hydrophobic patches alike.

Although ITC and MD are performed under different ionic environments, they prove complementary aspects of the same LLPS process. ITC experiments, conducted without added salt, quantify the intrinsic bulk behavior driven by the native charge complementarity of V_2_EV_2_ and V_2_KV_2_. In contrast, atomistic MD simulations necessarily include Na^+^ and Cl^-^ to maintain electrostatic neutrality under periodic boundary conditions, providing access to the nanoscale mechanisms that cannot be resolved calorimetrically. Crucially, the asymmetric association detected by ITC is fully reproduced in the simulations, despite electrostatic screening by dissolved ions. Spontaneous condensation in silico — featuring selective counterion release, solvation-shell depletion, chain compaction, and persistent differences in effective charge exposure— demonstrates that the thermodynamic driving forces are not abolished under saline stabilization. Instead, these results reinforce that condensation is primarily entropy-driven, arising from cooperative water and counterion release and the collective loss of solvation constraints. The convergence of ITC and MD —bulk thermodynamics without ions and molecular reorganization in a neutral ionic environment— reveals a robust condensation regime, governed by sequence-encoded solvation asymmetries that dictate stoichiometry and the energetic landscape of condensate formation.

Importantly, the number of interchain hydrogen bonds between V_2_EV_2_ and V_2_KV_2_ increased sharply after condensation. This enhancement in hydrogen bonding parallels the steep endothermic transitions observed in ITC. These interactions might contribute to stabilizing the condensed state but likely act in concert with the entropic gain from water and counterion release. In parallel, intramolecular hydrogen bonds become more stable and frequent above the *T*_t_, particularly within V_2_EV_2_, reinforcing its tendency to adopt globular conformations. Radial distribution functions (RDF) between charged residues and ions revealed a clear asymmetry in ion-protein interactions. Na^+^ ions preferentially associate with the negatively charged carboxylates of V_2_EV_2_, whereas Cl⁻ ions accumulate around the protonated ammonium groups of V_2_KV_2_. This differential ion coordination establishes distinct solvation shells and generates an unequal electrostatic potential across the two polypeptides. As a result, V_2_EV_2_ remains more tightly hydrated and locally screened, while V_2_KV_2_ retains a higher effective positive valence due to partial exposure of its cationic side chains. These solvation asymmetries modulate the electrostatic landscape of the system and directly influence its binding stoichiometry, explaining why an excess of V_2_KV_2_ is required to reach charge compensation in the condensate.

Together, calorimetric and atomistic simulation data provide a coherent, multiscale perspective on the thermoresponsive LLPS of V_2_EV_2_ and V_2_KV_2_. ITC revealed a high-affinity, enthalpically driven, yet asymmetric assembly process above the *T*_t_, requiring excess of V_2_KV_2_. This compositional asymmetry aligns closely with structural features extracted from the MD simulations. Snapshots (**Figure 3e, f**) demonstrate a clear transition from a dispersed, solvated state to a compact, aggregated configuration as temperature increases. In particular, clustering of V_2_EV_2_ and V_2_KV_2_ chains at 60*^◦^*C is accompanied by visible depletion of surrounding counterions, consistent with a release of solvation constraints—a process expected to contribute positively to the net entropy change upon complexation.

The progressive reduction in *R_g_* for both IDPPs, (**Figure 3g–h**), reinforces the calorimetric interpretation. The ∼4 °A decrease observed for both chains was temporally correlated with the formation of stable hydrogen-bonded contacts and reduced solvent accessibility, suggesting that the macromolecular compaction is not merely entropically favored, but also structurally stabilized.[47] This nanoscale structural reorganization is a hallmark of the phase separation and confirms that the calorimetric signals captured in ITC reflect real molecular rearrangements within the condensate.

### 2.4. pH–driven LLPS reveals opposite responses in charged IDPP condensates

To examine whether dense-phase chemistry can flip phase equilibria, we quantified apparent pK_aE_ values across temperatures below and above *T*_t_. We compared two pH-responsive condensate systems that behave oppositely to the same pH stimulus: V_2_EV_2_ and V_2_KV_2_-V_2_EV_2_ (**Figure 3**) to determine whether changes in local dielectric and ion partitioning within the condensate renormalize acid–base energetics, altering the sign and magnitude of ΔH°/ΔS° and even the temperature dependence of pK_aE_. In doing so, we establish whether protonation/deprotonation acts as a molecular switch that either disrupts (V_2_EV_2_) or stabilizes (V_2_KV_2_-V_2_EV_2_) condensation.

Acid-base titrations provide direct information on the apparent pK_aE_ of Glu residues under different local environments (i.e., fully solvated in dilute solution or embedded within a dense phase), thereby allowing the extraction of thermodynamic parameters (ΔH° and ΔS°) of the ionization process from Van’t Hoff analysis (**Figure 3i, j**).

Below the *T*_t_, both systems remained soluble and showed systematic increase in pK_aE_ with temperature (**Table S3**) with ΔH°<0, ΔS°<0 (**Table S4**). The formation of the carboxylate is exothermic because hydration of the anion and association of counterions release heat, yet it is entropically penalized by ordering of water and ions. The rise of pK_aE_ with temperature follows naturally as the dielectric constant of water decreases and dehydration of the charged state becomes more costly, making the acid effectively weaker in bulk solution.

This is consistent with strong hydration of the carboxylate anions and counterion binding in bulk solution.[41] The increase of pK_aE_ reflects the reduced stabilization of the deprotonated state at higher temperatures, as the dielectric constant of water decreases and dehydration becomes more costly, rendering the acid effectively weaker.

Above the *T*_t_, the two systems diverged. On the one hand, V_2_EV_2_ maintained the same trend (i.e., pK_aE_ increased with temperature and ΔH°<0, ΔS°<0), and the condensate dissolved by deprotonation (**Figure 3b**). Mechanistically, ionization breaks intermolecular contacts within the dense state and rehydrates exposed carboxylates, adding energetic and entropic penalties relative to the soluble regime. Importantly, although the sign of ΔH° and ΔS° remains negative, their absolute values are larger above *T*_t_ than below it (**Table S4**). This reflects the extra energetic cost and entropic penalty associated with breaking intermolecular contacts inside the condensate and rehydrating side chains, together with the restructuring of the solvent shell around newly exposed carboxylates and counterions.

On the other hand, V_2_KV_2_-V_2_EV_2_ exhibited an inverted response above the *T*_t_. Its apparent pK_aE_ decreased with increasing temperature (**Table S3**), and both ΔH° and ΔS° become positive (**Table S4**). Here, deprotonation is endothermic (enthalpically costly) but entropically favorable, indicating that the release of water and counterions and the formation of stabilizing salt bridges dominates the free-energy balance (**Figure 3f,h**). At acidic pH (pH < pK_aE_), the Glu residues remain protonated, and therefore the V_2_EV_2_ blocks tend to collapse. However, the V_2_KV_2_-V_2_EV_2_ remained soluble by the highly solvated cationic V_2_KV_2_ block, which prevents condensation. As the pH increased (pH > pK_aE_), deprotonated Glu-COO^-^ engaged in electrostatic pairing with Lys-NH_3_^+^ residues, promoting salt-bridge formation, water release, and ultimately hydrophobic collapse of the diblock copolymer. The large positive enthalpy of deprotonation for V_2_KV_2_-V_2_EV_2_ above the *T*_t_ (ΔH° = 14.9 kcal·mol^-1^) compared to the negative enthalpy measured for V_2_EV_2_ under the same conditions (ΔH° = -6.7 kcal·mol^-1^) reflects their opposite condensation behavior. In V_2_KV_2_-V_2_EV_2_, deprotonation promotes condensation, whereas in V_2_EV_2_ it induces solubilization. Although chain length contributes to the total enthalpy (V_2_KV_2_-V_2_EV_2_ contains twice as many pentapeptide repeats) the magnitude of ΔH° arises primarily from the cooperative reorganization of its diblock architecture. The simultaneous formation of salt-bridge networks and enhanced hydrophobic contacts amplifies the energetic cost of ionization while stabilizing the condensed state, as corroborated by the stronger reversing signal observed in TM-DSC (**Figure 2f**).

Once two phases coexist in charged condensate systems, ions distribute unequally between the dense condensate and dilute phases.[48,49] This unequal ion partitioning might create an electrostatic potential difference between the two phases, which defines a distinct internal pH and ion activity inside the condensate. Because the apparent pK_aE_ of ionizable residues is referenced to the external pH, this internal potential effectively shifts the measured pK_aE_ values within the dense phase and can even invert the temperature dependence observed for V_2_EV_2_ and V_2_KV_2_-V_2_EV_2_. In parallel, changes in the microenvironment of the condensate, i.e., changes in effective dielectric constant, hydration, and viscoelastic network,[30,50] also influences which ionization state is stabilized.[48,50]

These combined effects delineate two pH-responsive routes to condensation. V_2_EV_2_ exemplifies a protonation/neutralization pathway, where acidification reduces electrostatic repulsion, increases backbone proximity, and promotes water release, stabilizing condensed state. This mechanism simulates the behavior of native acidic IDPs, such as RPT segment of Pmel17,[51] which is soluble at cytosolic pH yet solidifies into functional amyloid upon melanosomal acidification. In that system, the emergence of intermolecular charge-transfer transitions during aging reflects progressive chain compactation and water release. [52] In contrast, V_2_KV_2_-V_2_EV_2_ exemplifies a deprotonation/ion-pairing pathway. Above the *T*_t_, a cation-rich, lower-dielectric interior favors Glu⁻-Lys⁺ pairing, thus lowering the dense-phase pK_aE_, and reinforces LLPS despite the endothermic cost of ionization.

The opposite pK_aE_(T) trends and the signs and magnitudes of ΔH°/ΔS° obtained for V_2_EV_2_ versus V_2_KV_2_-V_2_EV_2_ align precisely with this model. In V_2_EV_2_, deprotonation is exothermic and entropically penalized (ΔH° < 0, ΔS° < 0), reflecting hydration-dominated energetics that drive condensate dissolution. In V_2_KV_2_-V_2_EV_2_, deprotonation becomes endothermic yet entropically favored (ΔH° > 0, ΔS° > 0), with a lower apparent pK_a_ above the *T*_t_. This behavior arises because salt-bridge formation and the release of counterions and bound water within a low-dielectric, cation-rich microenvironment stabilize the charged state and reinforce phase separation.

### 2.5. Intramolecular salt-bridges encode condensate microenvironment

To connect salt-bridge topology to the condensate microenvironment and miscibility, we combined NMR reporters of Lys protonation with confocal microscopy to visualize mixing behavior in the presence of a non-charged partner (V). NMR resolves the fraction of inter- vs. intramolecular salt bridges, a topological descriptor that sets residual free charge in the dense phase. By correlating this topology with partitioning behavior and condensate architecture, we tested whether sequence-encoded ionic connectivity governs micropolarity, miscibility, and the transition between single-phase and multiphase condensates.

Direct detection of salt-bridge formation by 2D-NMR (e.g., ^1^H–^15^N HMBC and ^1^H-^1^H NOESY) was attempted but proved not feasible due to severe signal broadening in the condensed phase (**Figure S8-S15**). This broadening reflects the limitations of the technique in highly crowded protein phases, where restricted molecular motion and rapid transverse relaxation (T_2_) suppress long-range correlations and exacerbate peak overlap. Although occasional weak cross-peaks were observed, they could not be unequivocally assigned to specific interactions between Glu and Lys side chains, impeding precise quantification of these contacts (**Figure S14**). Instead, we monitored the ε-methylene hydrogens of Lys, adjacent to the terminal primary amine, which serve as sensitive reporters for its effective protonation state (**Figure 5a**). At pH 7, these hydrogens produced a resonance at 2.88 ppm and, upon increasing pH, the signal shifted progressively upfield consistent with deprotonation of the Lys ammonium group and a reduction of the local positive charge (**Figure 5b**).

**Figure 5.**
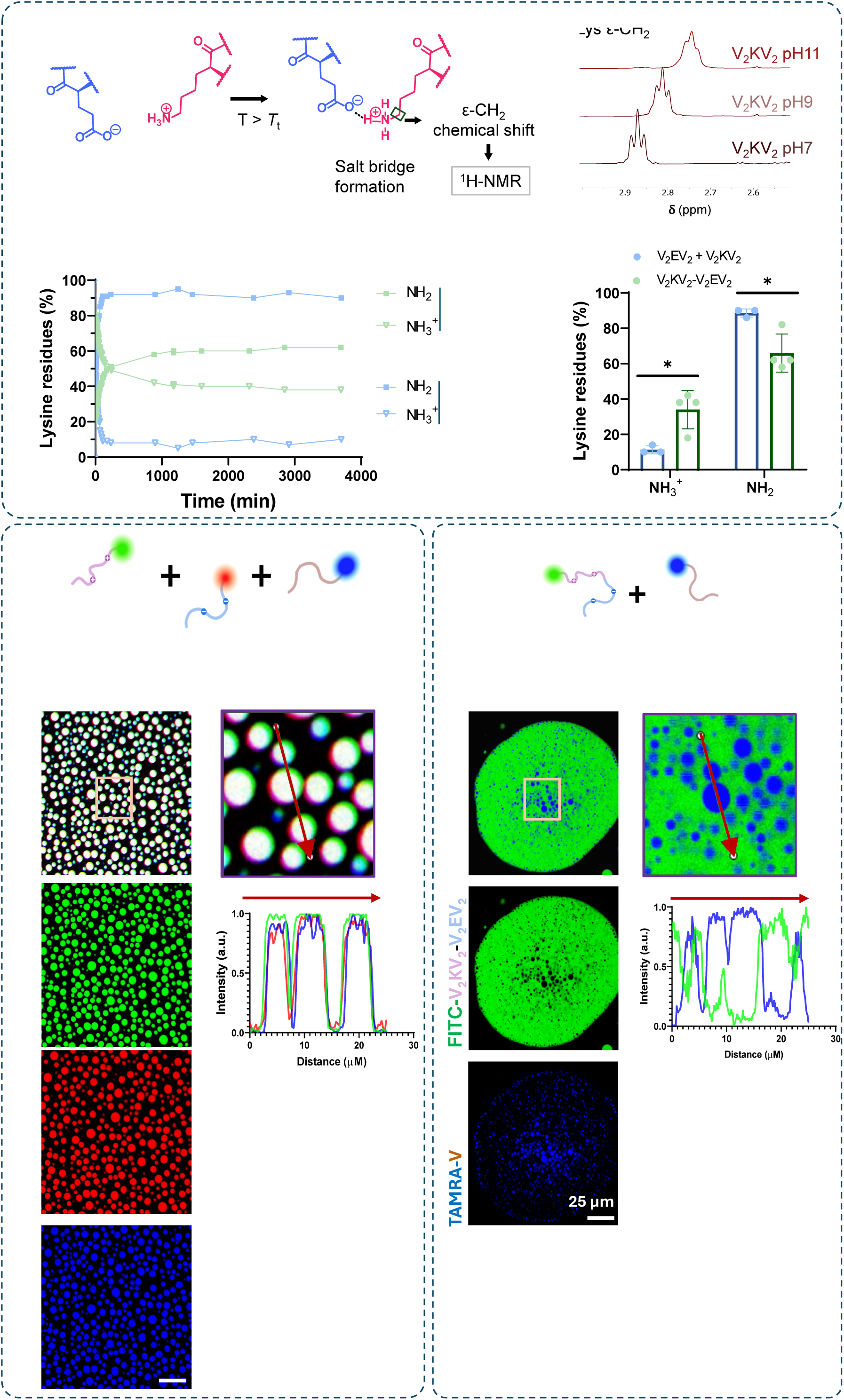
Salt-bridge formation modulates internal polarity and partitioning behavior in IDPP condensates. a) Schematic representation of electrostatic pairing between glutamic acid and lysine side chains, highlighting salt-bridge formation through carboxylate-ammonium interactions. b) Representative ^1^H-NMR spectra of V_2_KV_2_ at pH 7, 9, and 11. The chemical shift of the ε-methylene protons (-CH_2_-) adjacent to the terminal amine group (Lys) shift upfield with increasing pH, consistent with reduced local positive charge and Lys deprotonation. This signal was used as a reporter of the protonation state during salt-bridge formation. c) Time-resolved quantification of NH_3_^+^ and NH_2_ species by NMR in V_2_KV_2_ + V_2_EV_2_ and V_2_KV_2_ – V_2_EV_2_ mixtures during LLPS. d) Final NH₂/NH₃⁺ ratios after 3000 min. V_2_EV_2_ + V_2_KV_2_ shows a significantly higher degree of deprotonation, indicating enhanced intramolecular salt-bridge formation. (e–f) Confocal microscopy images of condensates containing a third, neutral hydrophobic IDPP (V), mixed either with V_2_KV_2_ + V_2_EV_2_ (e) or with V_2_KV -V_2_EV_2_ (f). In V_2_KV_2_ + V_2_EV_2_, all components colocalize uniformly throughout the condensates. In contrast, in V_2_KV_2_-V_2_EV_2_ condensates, changes in polarity lead to the formation of discrete immiscible V-based microdomains. Fluorescence intensity profiles along representative condensates confirm distinct partitioning behaviors. Note: TAMRA is artificially displayed in blue to improve contrast.

A similar upfield displacement of the Lys ε-methylene resonance was observed for V_2_KV_2_ + V_2_EV_2_ mixtures driven above the *T*_t_ in D_2_O, even under controlled bulk-pH conditions (**Figure S16**). Because the external pH remains constant, this shift cannot result from simple basification of the medium. We interpret this effect as consistent with salt-bridge formation, whereby one proton from Lys–NH₃⁺ engages in electrostatic pairing with Glu–COO⁻, reducing the effective positive charge on the Lys–NH₃⁺ and thereby increasing the upshielding of the adjacent ε-methylene protons. This effect reproduces the spectral behavior observed upon basification in bulk solution but originates from charge delocalization within the condensate.

This approach enabled us to quantify the fraction of Lys residues involved in salt bridges (**Figure 5c, d**). In equimolar V_2_EV_2_+ V_2_KV_2_ mixtures, up to 90% of lysine residues participated, whereas in the diblock V_2_KV_2_-V_2_EV_2_ this fraction dropped to ∼70%. We attribute this reduction to intramolecular salt bridges within the diblock, which constrains conformational freedom and limits intermolecular pairing.

These measurements indicate that molecular architecture (mixture vs. diblock) determines both the density and topology (intra- vs intermolecular) of salt bridges, and therefore, the amount of residual free charges in the condensed phase. Because the condensate microenvironment governs condensate architecture, miscibility, and client partitioning,[30] we hypothesized that lower salt-bridge density, i.e., more free charges, would increase micropolarity and thus alter miscibility with other IDPs. To test this, we designed multicomponent condensates by combining the charged systems with a hydrophobic LCST-IDPP, V. Throughout this study, topology refers not to classical secondary/tertiary structure, but to the network connectivity of ionic contacts enforced by sequence architecture.

In the presence of IDPP V, both systems (equimolar V_2_EV_2_ + V_2_KV_2_ and covalently linked V_2_KV_2_-V_2_EV_2_) underwent LLPS, concentrating virtually all proteins into dense phases (**Figure 5e, f**). However, these synthetic condensates diverged markedly. In V_2_EV_2_ + V_2_KV_2_ + V mixtures, where inter-chain pairing is high (∼90%) and residual free charge is minimized, all three proteins co-assembled into homogeneous spherical condensates, with uniform distribution throughout the interior (**Figure 5e**). This is consistent with a lower-polarity interior compatible with the hydrophobic partner. By contrast, in V_2_KV_2_-V_2_EV_2_ + V mixtures, where intra-chain pairing is increased (∼70% total pairing) and more free charge is retained, V segregated into distinct internal compartments within the V_2_KV_2_-V_2_EV_2_ condensates (**Figure 5f**), consistent with a micropolarity mismatch that drives multiphase organization.

In this way, these findings establish a direct connection between protein polymer architecture, salt-bridge topology, and condensate polarity. High inter-chain pairing (V_2_EV_2_ + V_2_KV_2_) yields lower-polarity, compositionally homogeneous multicomponent condensates, whereas increased intra-chain pairing (V_2_KV_2_-V_2_EV_2_) preserves free charge and raises micropolarity, promoting internal demixing and multiphase condensation in the presence of a non-charged IDP partner. By biasing salt-bridge formation within a chain (intramolecular) or between chains (intermolecular), the sequence fixes the level of residual charge in the condensate. This, in turn, encodes dense-phase micropolarity and dictates phase architecture, promoting either single-phase organization or multiphase partitioning when mixed with additional partners. This architectural control of condensate polarity opens a route for bottom-up biofabrication. By tuning charge patterning, block arrangement, and stoichiometry, the same protein building blocks can generate condensates with programmable compartmentalization. We expect this protein system can be used to co-assemble with enzymes, cofactors, nucleic acids, and small-molecule substrates to create, for example, selectively partitioning microreactors that support compartmentalized catalysis, stimuli-responsive gating, and dynamic and tunable scaffolds for tissue engineering.[53,54] Such control on micropolarity using the same protein components also enables hierarchical condensate assemblies, reminiscent of the nucleolus in cells and consistent with in-vitro polymer systems,[34,55,56] thereby creating microenvironments with defined micropolarities and thus, biochemical properties. Moreover, the intrinsically charged nature of these condensates means that co-assembly with ions and small molecules offer a second way of control over dense-phase properties.[50,57] Hence, future work will focus on, systematically varying ionic identity, strength, and cargos of clinical relevance to reveal how these interactions tunes internal organization, micropolarity gradients and functionality.

## 3. Conclusion

In summary, we establish a sequence-encoded framework for programming the phase behavior, thermodynamic landscape, and microenvironmental properties of synthetic biomolecular condensates using minimalistic, genetically encoded LCST-IDPPs. By systematically varying the identity and density of charged residues together with molecular architecture, and stoichiometry, we reveal how the interplay between hydrophobic collapse and electrostatic interactions governs the onset, cooperativity, and internal organization of bicomponent IDPP condensates. Through an integrated approach combining calorimetry, spectroscopy, and molecular simulations, we uncover two distinct energetic regimes (i.e., slow cooperative dehydration and rapid chain reorganization) that emerge from the interplay between intra- and intermolecular salt-bridge formation, solvent reorganization, and chain desolvation. The results reveal that the topology of ionic interactions, whether encoded as interchain or intrachain pairings, determines the residual charge and micropolarity of the dense phase and thereby controls the miscibility of condensates with other partners. These findings provide a quantitative and mechanistic framework to rationally design synthetic condensates with programmable phase behavior and internal polarity, bridging the gap between reductionist IDPP models and the complex multiphase organization of natural biomolecular condensates. Beyond their fundamental relevance, these insights open new avenues for engineering responsive microenvironments, selective molecular partitioning, and dynamic scaffolds for catalysis, biosensing and tissue engineering.

## CRediT authorship contribution statement

**Julio Fernandez-Fernandez:** Visualization, Validation, Methodology, Investigation, Formal analysis. **Vicente Dominguez Arca:** Writing – review & editing, Visualization, Validation, Investigation, Formal analysis, Data curation. **Raul Escribano:** Visualization, Validation, Investigation, Formal analysis. **Sergio Ferrero:** Writing – review & editing, Visualization, Validation, Investigation, Formal analysis, Data curation. **Sergio Acosta:** Writing – original draft, review & editing, Methodology, Visualization, Investigation, Formal analysis, Data curation, Supervision, Conceptualization. **José Carlos Rodríguez-Cabello:** Writing – review & editing, Visualization, Methodology, Supervision, Funding acquisition, Conceptualization.

## Acknowledgements

This work was funded by Spanish Government (grants PID2021-122444OB-100 and PID2022-137484OB-I00 funded by MCIN/AEI/ 10.13039/501100011033 and by ERDF, EU), the *Junta de Castilla y León* (grant VA188P23 and CLU-2023-1-05 cofunded by ERDF), and *Centro en Red de Medicina Regenerativa y Terapia Celular de Castilla y León*. V.D.A. acknowledges financial support from *Xunta de Galicia* (grant IN606B-2023/006). The authors thank Viktoriya Chaskovska for assistance with cloning the encoding genes into the expression plasmids, Rocío García Lera for support with IDPP production and purification, and Dr. Javier Reguera for helpful discussions.

## Use of artificial intelligence tools

During the preparation of this work the author used ChatGPT exclusively to refine and modify phrasing within the manuscript. After using this tool, the authors reviewed and edited the content as needed and takes full responsibility for the content of the publication.

