## Supplementary Information for "Topology-Encoded Polarity in Oppositely Charged Binary IDPP Condensates: Multiphase Organization from Non-Coacervating Partners as a Minimal Model of Complex Coacervation"

**Materials and Methods**

**Production and characterization of the IDPP library**

The IDPPs were produced recombinantly in *Escherichia coli* following previously described methods. [1] Briefly, the encoding genes were purchased from NZYTech (Portugal) in the pUC57 cloning plasmid, subcloned into an expression vector derived from pET25(+), and transformed into *E. coli* BLR(DE3).

Recombinant expression was carried out by bacterial fermentation in a 15 L bioreactor (Applikon) using modified Terrific Broth (TB) medium under continuous stirring at 500 rpm and 37 °C. After 18 h of fermentation, the cells were harvested by centrifugation at 4 °C and washed with isotonic buffer to remove residual fermentation components. The cell pellets were then lysed by disruption (GEA Panda 2000).

Purification of the IDPPs was achieved through their lower critical solution temperature (LCST) phase behavior using inverse transition cycling (ITC). [2] The purified proteins were subsequently dissolved and dialyzed extensively against 25 L of ultrapure water (type II, three changes and type I, one change) at 4 °C, sterilized by filtration (0.22 µm, Nalgene), and lyophilized (Labconco). Typical yields ranged from 530 to 750 mg of purified polymer per litre of culture medium.

The monodispersity and purity of the IDPPs were evaluated by sodiumdodecyl sulfate-polyacrylamide gel electrophoresis (SDS-PAGE), matrix-assisted laser desorption/ionization time-of-flight (MALDI-ToF) mass spectrometry, and high-performance liquid chromatography (HPLC), and nuclear magnetic resonance (NMR). The MALDI-ToF, HPLC and NMR experiments were carried out at Laboratory of Instrumental Techniques (LTI) Research Facilities, University of Valladolid.

**Chemical functionalization for confocal microscopy**

Fluorescent derivatives of the IDPP library were prepared by covalent conjugation of fluorescent probes, linked N-hydroxysuccinimide (NHS), to primary amines via nucleophilic substitution. Briefly, solutions of NHS-dyes (i.e., Alexa Fluor 647 NHS ester, fluorescein isothiocyanate (FITC), and 5(6)-carboxytetramethylrhodamine N-succinimidyl ester (TAMRA-SE) (all from Thermo Fisher Scientific)) were dissolved in anhydrous DMSO and mixed with IDPP solutions in anhydrous DMF at a 1:1 molar ratio (dye:IDPP). The reaction mixtures were stirred for 24 h at room temperature under a nitrogen atmosphere.

After completion, the products were dialyzed repeatedly against 5L of ultrapure water (three changes in type II and two changes in type I) at 5 °C, sterilized by filtration through 0.22 μm membranes, and lyophilized.

For all confocal microscopy experiments, the fluorescently labelled IDPPs were mixed with their unlabeled counterparts to ensure that 10 mol% of the total polymer in solution was labelled, thereby minimizing perturbation of the native phase-separation behavior.

**Condensate visualization setup**

For visualization, 70 μL of 250 μM of IDPP solution was placed on a glass slide (Fisherbrand™, 12373118) equipped with an incubation chamber composed of an adhesive spacer frame (Frame-Seal BIO-RAD, SLF0601) and a cover glass (Epredia™, 15747592) positioned on top of the spacer.

Both the slide and the cover glass were subjected to a Pluronic F-127 (Sigma Aldrich) treatment to prevent surface adhesion, creating a non-stick layer on the glass surfaces to inhibit the attachment of the IDPPs.[3] This treatment consisted of cleaning with 70% (v/v) ethanol, followed by rinsing with ultrapure (type I) water and air-drying under a laminar flow hood. The components were then immersed in a freshly prepared 5% (v/v) Pluronic F-127 solution for one hour at room temperature. Unbound Pluronic was subsequently removed by washing ten times (1 min per wash) with ultrapure water. Finally, the treated components were air-dried in a sterile environment before use.

**Confocal microscopy**

Fluorescence images were captured by TCS SP8 confocal microscope (Leica Microsystems, Germany) equipped with a 405-nm laser and a pulsed white light laser (WLL). Imaging was performed with a 40x oil-immersion objective (HC PL Apo 40x/1.30oil CS2) integrated with an environmental heating chamber for temperature control. Excitation wavelengths were set to 491 nm for FITC, 552 nm for TAMRA, and 653 nm for Alexa Fluor 647.

Prior to imaging, the IDPP solutions were incubated within the sealed cover-glass assembly at 60 °C for 20 min to induce liquid–liquid phase separation (LLPS). Image acquisition and processing were performed using Leica Application Suite X (LAS X) software.

**Evaluation of the thermal behavior by turbidimetry**

Turbidimetry experiments were conducted using a Varian Cary 100 UV-Vis spectrophotometer (Varian Inc., NC, USA) equipped with a temperature-controlled cuvette holder. The optical density at 350 nm ($OD^{350}$) of the IDPP solutions was recorded different conditions.

Two sets of experiments were carried out both in ultrapure water type I and in PBS buffer. In the first set, a concentration series of IDPPs, including a physical mixture of $V_2EV_2$ and $V_2KV_2$ ($V_2EV_2$ + $V_2KV_2$), was prepared at 10, 15, 25, 50, 75, 100, 150, 200, 300, 500, and 1000 µM. In the second set, the influence of charge stoichiometry on coacervation was investigated by mixing V4E1 and V4K1 at different volumetric ratios (100:0, 80:20, 60:40, 50:50, 40:60, 20:80, 0:100) at a fixed final concentration of each protein component of 25 µM.

All data were collected at a scan rate of 1 °C $min^{-1}$, and each condition was analysed in triplicate. The transition (cloud point) temperature ($T_t$) was determined as the temperature corresponding to the maximum of the first derivative of the $OD^{350}$ vs temperature curve.

**Enthalpy analysis by Differential Scanning Calorimetry (DSC) and Temperature-Modulated DSC (TM-DSC)**

DSC and TM-DSC experiments were performed using a Mettler Toledo 822e calorimeter equipped with a liquid nitrogen cooling system. IDPP solutions (50 mg $mL^{-1}$) were prepared in ultrapure water type I at different pH values and in PBS buffer. In addition, mixtures of $V_2EV_2$ and $V_2KV_2$ at the same volumetric ratios used in the turbidimetry experiments were analyzed under identical conditions. For all experiments, 20 µL of prepared solutions were introduced into a standard 40 µL aluminium pan, which were hermetically sealed. An equivalent volume of solvent was added to a reference pan.

In conventional DSC experiments, samples were equilibrated for 2 min at 20 °C and subsequently heated at a constant rate of 2 °C $min^{-1}$. TM-DSC measurements were conducted by superimposing a sinusoidal temperature modulation onto a linear heating ramp. The modulation parameters were: underlying heating rate (ν) = 1 °C $min^{-1}$, amplitude (A) = 0.1 °C, and period (P) = 0.6 min. The selected TM-DSC conditions correspond to those previously optimized for the analysis of this class of IDPPs, ensuring reliable deconvolution of the reversing and non-reversing enthalpic components. [4,5]

**Evaluation of the secondary structures by circular dichroism (CD) spectroscopy**

The secondary structures of IDPP library and mixtures from the $V_2EV_2$ + $V_2KV_2$ system were analyzed by CD spectroscopy using a Jasco J-810 spectropolarimeter (Jasco, U.S.A.) equipped with 0.1 cm quartz cells and with a temperature controller (STTI, University of Alicante, Spain).

Samples were prepared at a concentration of 0.35 mg $mL^{-1}$ in ultrapure water, and the pH was adjusted to 7. Experiments were conducted below and above the $T_t$, specifically at 20 ºC and 60 ºC, with a stabilisation time of 20 minutes prior to data acquisition. CD spectra were recorded in the wavelength range of 190-250 nm. A 15 pt Savitzky-Golay filter smoothed data.

The secondary-structure content of each sample was estimated from the experimental spectra using the BeStSel algorithm,[6,7] which enables quantitative deconvolution of β-sheet, α-helix, and disordered contributions in proteins.

**Analysis of the thermodynamic parameters of interactions by isothermal Titration Calorimetry (ITC)**

The binding affinity and stoichiometry governing co-assembly between $V_2EV_2$ and $V_2KV_2$ were determined below (22 °C) and above (60 °C) the $T_t$ using a MicroCal VP-ITC microcalorimeter (Malvern) at the Soft Materials Service – U6 NANBIOSIS, Institute of Materials Science of Barcelona (CSIC, Spain).

To probe the directionality of binding, two titration schemes were conducted at each temperature: (1) $V_2EV_2$ in the calorimetric cell and $V_2KV_2$ in the syringe, and (2) $V_2KV_2$ in the cell and $V_2EV_2$

in the syringe. Both IDPPs were dissolved in ultrapure water to minimize baseline heat from buffer mismatch. The concentration in the cell was 57.9 µM, and in the syringe 579.6 µM. Each titration consisted of 35 injections of 9 µL. Control titrations ($V_2KV_2$ or $V_2EV_2$ into ultrapure water) were performed to account for dilution heats, which were subtracted from the raw data to obtain the net heat of binding.

The theoretical model used for fitting is based on the standard equilibrium binding formalism. The fraction of occupied binding sites, θ, is defined in terms of the equilibrium constant K and the free ligand concentration ([X]) by the equation:

$$K = \frac{\theta}{(1-\theta)[X]}$$

This relationship is coupled with the conservation of mass, which for a macromolecule of total concentration Mt and a ligand of total concentration Xt, takes the form:

$$Xt = [X] + n\theta Mt$$

Combining the two expressions yields a quadratic equation in θ :

$$\theta^2 - \theta\left(1 + \frac{Xt}{nMt} + \frac{1}{nKMt}\right) + \frac{Xt}{nMt} = 0$$

Once θ is solved at each injection point, the total heat content Q of the cell at fractional saturation is calculated as:

$$Q = nMt\Delta HV0\theta$$

where ΔH is the molar binding enthalpy and V0 is the cell volume. However, since titration involves the addition of finite aliquots of ligand solution, a correction must be applied for the displaced volume after each injection. This correction accounts for the fact that a portion of the solution previously in the cell is ejected with each addition, yet still contributes to the observed heat signal due to fast mixing and reaction kinetics. The corrected heat evolved after the ith injection, ΔQ(i), is then computed as:

$$\Delta Q(i) = Q(i) + \frac{\delta Vi}{V0}\left[\frac{Q(i) + Q(i-1)}{2}\right] - Q(i-1)$$

This expression was applied iteratively during nonlinear regression fitting, using a Marquardt-Levenberg algorithm to minimize the residuals between experimental and calculated heats. The parameters of interest—stoichiometry n, binding constant K, and enthalpy ΔH—were extracted for both titration directions.

**Computational Methods**

Atomistic molecular dynamics (MD) simulations were conducted to resolve the nanoscale structural rearrangements underlying thermoresponsive coacervation in mixtures of V2EV2 and V2KV2. Initial chain conformations were generated using AlphaFold2, which predicted highly flexible structures with limited tertiary organization,[8] consistent with the expected behavior of IDPPs with LSCT behavior (i.e., ELRs). For $V_2EV_2$, the predicted structural metrics included ptm values 0.16–0.26 and fraction of disorder ~0.25 . $V_2KV_2$ exhibited ptm $\approx$ 0.24 and ~0.21 disorder, with low chain-pair confidence (chain_pair_iptm $\approx$ 0.24). These modest confidence values reflect the inherent conformational heterogeneity of disordered polypeptides and do not indicate structural inaccuracies. Instead, they validate that both ELRs exist as dynamic ensembles lacking rigid folded domains, a prerequisite for reversible coacervation.

The predicted structures were placed in a cubic periodic box of explicit water (TIP3P model), and $Na^+$ and $Cl^-$ ions were added to ensure overall charge neutrality and prevent unphysical intrachain electrostatic repulsion under periodic boundary conditions. An equimolar 1:1 mixture of $V_2EV_2$ and $V_2KV_2$ was simulated to probe the onset of intermolecular associations driving condensation.

All simulations employed the CHARMM36m all-atom force field, optimized for intrinsically disordered proteins, along with CHARMM parameters for monovalent ions. Long-range electrostatic interactions were evaluated using particle mesh Ewald (PME) with a 10–12 Å real-space cutoff. Bond constraints involving hydrogens were enforced with LINCS, allowing a 2 fs time step. Temperature and pressure were controlled using the Nosé–Hoover thermostat and Parrinello–Rahman barostat at 20°C (below the transition temperature) and 60°C (above Tt), and 1 bar, respectively.

All systems underwent energy minimization, followed by NVT equilibration (200 – 500 ps) and NPT equilibration (1 – 2 ns), prior to production runs of ⩾200 ns per temperature condition. Analyses were performed using GROMACS and VMD, quantifying the radius of gyration (Rg), solvent-accessible surface area (SASA), inter- and intrachain hydrogen bonding, and radial distribution functions (RDFs) between charged residues and counterions. Temporal correlations of these features enabled evaluation of thermally induced compaction, hydration-layer depletion, and ion redistribution within emerging protein-rich clusters.

These MD simulations were not intended to reproduce absolute calorimetric values —ITC experiments were performed without added buffer salts— but instead to provide a mechanistic interpretation of the entropy-driven coacervation detected experimentally. The emergence of compositional asymmetry and spontaneous interchain association even under electrostatic screening confirms that counterion and water release, rather than mere Coulombic pairing, dominate the driving forces behind condensate formation in this two-component ELR system.

**Determination of apparent $pK_a$ values by acid–base titration**

Acid–base titrations were performed using a TitroLine 7000 automatic titrator (SI Analytics, Germany) equipped with a 20 mL burette and an A162 2M-DIN combined glass electrode. The sample temperature was controlled by connecting a refrigerated circulating bath to a double-jacketed reaction flask (Merck), ensuring continuous water flow through the system during measurements.

Before each experiment, the pH electrode was calibrated at the measurement temperature using standard buffer solutions. A three-point calibration was automatically executed by the instrument to guarantee precision and stability across the full titration range.

For each titration, samples were dissolved in 5 mL of ultrapure water to a final IDPP concentration of 2.5 mM. The sample mass was adjusted to maintain an equivalent total number of carboxylic groups across all systems, allowing direct comparison of ionizable site behavior. The initial pH was adjusted to approximately 2.0 by incremental addition of 0.1 M HCl.

Titrations were carried out under continuous magnetic stirring at 500 rpm using the instrument's control software. A 55 mM NaOH solution was delivered from the burette at a constant flow rate of 50 $\mu L \cdot min^{-1}$ until the pH reached approximately 11.0. The pH and titrant volume were recorded in real time throughout the titration.

Apparent $pK_a$ values were extracted from the titration profiles as the midpoint of the buffer region corresponding to the maximum of the first derivative (dpH/dV). When the inflection point was not sharply defined, $pK_a$ values were determined by linear interpolation of the pre- and post-inflection baselines to minimize experimental uncertainty.

**Characterization by nuclear magnetic resonance (NMR)**

NMR experiments at various temperatures were measured by 500 MHZ Agilent DD2 instruments equipped with a cold probe in the Laboratory of Instrumental Techniques (LTI) Research Facilities, University of Valladolid. Chemical shifts for $^1H$, $^{13}C$, $^{15}N$ nuclei were measured in ppm, with the HDO residual solvent signal used as an internal reference or nitromethane ($CH_3NO_2$).

Two different temperatures (25º and 60 ºC) were employed during the 2D NMR studies. Classical 2D NMR methods were employed in order to achieve $^1H$, $^{13}C$ and $^{15}N$ peak assignments employing 500 MHZ Agilent DD2 instruments equipped with a HFX indirect probe or oneNMR probe in the Laboratory of Instrumental Techniques (LTI) Research Facilities, University of Valladolid. For heteronuclei such as carbon-13 and nitrogen-15, frequency-swept (adiabatic) pulses in place of hard 180-degree pulses were used to improve spectral quality.

Homonuclear $^1H$–$^1H$ 2D experiments total correlation spectroscopy (zTOCSY) and nuclear Overhauser effect spectroscopy (NOESY) with a zero quantum filter were used for artifact suppression.[9] zTOCSY experiments were acquired in the phase-sensitive mode using the PRESAT pulse sequence in order to suppress the residual water signal resonance. A total of four transients for each of the 200 t1 increments were recorded, using a spectral width of 2323.42 Hz for both dimensions (a total of 12 Hz resolution in indirect dimension), with a mixing time for the DIPSI2 spin lock of 100 ms and an acquisition time of 150 ms. The data were apodized with Gaussian window in both dimensions. NOESY experiments were performed using the PRESAT pulse sequence to remove the residual water signal resonance, with the following acquisition parameters: 16 transients for each of the 400 t1 increments, 2323.42 Hz spectra width for both dimension (a total of 6 Hz resolution in indirect dimension), and an acquisition time of 150 ms. A short vale of mixing times (200 ms) was used to avoid dreadful spin diffusion problems.[10] Data were fitted with Gaussian window in both dimensions. Carbon signals were measured indirectly using $^1H$-$^{13}C$ HSQC and HMQC experiments. They were acquired with inverse detention and carbon decoupling during acquisition in the phase-sensitive mode, with PRESAT solvent suppression, employing the following acquisition parameters: 64 transients for each of the 128 t1 increments, 2323.42 Hz spectral width in F2 dimension and 8547.12 Hz in the F1 dimension (a total of 67 Hz resolution in indirect dimension) and 146 Hz as a nominal value for the one-bond coupling constant $J_{CH}$. Data were fitted with Gaussian window in both dimensions. Nitrogen signals were obtained indirectly using $^1H$-$^{15}N$ HMBC sequence in the phase-sensitive mode with PRESAT solvent suppression, employing the following acquisition parameters: 512 or 800 transients for each of the 30 t1 increments, 2385.5 Hz spectral width in F2 dimension and 12158.1 Hz in the F1 dimension (a total of 405 Hz resolution in 15N indirect dimension) and 3 or 5 Hz as a nominal value for the long-range coupling constant nJNH. Data were fitted with sinebell window in F2 dimension and Gaussian window in F1 dimension.

All spectra were post-processed and analysed using Mestrelab Research software (MNova 15.0) and VnmrJ4.2 software (Agilent)

**Statistical analysis**

All experiments were performed with at three independent replicates unless otherwise stated. Data are presented as mean ± standard deviation (SD). Statistical analyses were performed using GraphPad Prism 8. For calorimetric analyses, sample sizes were as follows: TM-DSC (n = 3); DSC in ultrapure water: V (n = 7), V–V (n = 5), $V_2KV_2$-$V_2EV_2$ (n = 6), $V_2EV_2$ + $V_2KV_2$ mixtures (n = 7), $V_2EV_2$ (n = 3), and $V_2KV_2$ (n = 3); DSC in PBS: V (n = 7), V–V (n = 7), $V_2KV_2$– $V_2EV_2$ (n = 6), $V_2EV_2$ + $V_2KV_2$ mixtures (n = 4–6, depending on the molar ratio), $V_2EV_2$ (n = 3), and $V_2KV_2$ (n = 3).One-way ANOVA followed by Tukey's multiple-comparison test was applied to CD spectroscopy data (BeStSel) to evaluate differences in secondary-structure content among conditions. For NMR experiments quantifying lysine protonation states, the percentage of deprotonated lysine residues (%$NH_2$) was analyzed by two-way ANOVA followed by Sidak's multiple-comparison test.

### Experimental data

**Table S1.** Amino-acid sequences, theoretical and experimental molecular weights, and isoelectric points of the IDPPs

| IDPP | Sequence (aa) | Theo. Mw[a] (Da) | Exp. Mw[b] (Da) |
|---|---|---|---|
| $V_2KV_2$ | MESLLP [(VPGVG)$_2$VPGKG(VPGVG)$_2$]$_{10}$V | 21552.67 | 21545.67 ± 0.58 |
| $V_2EV_2$ | MESLLP [(VPGVG)$_2$VPGEG(VPGVG)$_2$]$_{10}$V | 21562.08 | 21569.67 ± 8.50 |
| $V_2KV_2$- $V_2EV_2$ | MESLLP[(VPGVG)$_2$VPGKG(VPGVG)$_2$]$_{10}$[(VPGVG)$_2$VPGEG(VPGVG)$_2$]$_{10}$V | 42326.78 | 42349.00 ± 2.65 |
| V | MESLLP(VPGVG)$_{50}$V | 21262.25 | 21261.00 ± 1.73 |
| V-V | MESLLP(VPGVG)$_{100}$V | 41736.54 | 41590.33 ± 9.81 |

[a]The theoretical molecular weight (Theo. Mw ) was calculated with the ProtParam tool (https://web.expasy.org/protparam/). [b]The experimental Mw (Exp. Mw) was measured by mass spectroscopy (Fig. S2).

**Table S2.** Amino acid composition of the recombinant IDPP library determined by HPLC amino-acid analysis after acid hydrolysis.

For each construct theoretical residue counts (Theo) derived from the designed sequences are compared with experimental counts (Exp) obtained from purified samples. The close agreement between Theo and Exp across all constructs confirms the expected sequences and high sample purity.

| | | $V_2EV_2$ | | $V_2KV_2$ | | $V_2KV_2$- $V_2EV_2$ | | V | | V-V | |
|---|---|---|---|---|---|---|---|---|---|---|---|
| | AA | Theo | Exp | Theo | Exp | Theo | Exp | Theo | Exp | Theo | Exp |
| D | ASP | | | | | | | | | | |
| E | GLU | 11 | 11.83 | 1 | 1.28 | 11 | 8.56 | | | | |
| N | ASN | | | | | | | | | | |
| S | SER | 1 | 0.87 | 1 | 1.26 | 1 | 1.31 | 1 | 1.07 | 1 | 2.81 |
| Q | GLN | | | | | | | | | | |
| H | HIS | | | | | | | | | | |
| G | GLY | 100 | 103.94 | 100 | 102.06 | 200 | 199.07 | 100 | 99.32 | 100 | 1.41 |
| T | THR | | | | | | | | | | |
| R | ARG | | | | | | | | | | |
| A | ALA | | | | | | | | | | |
| Y | TYR | | | | | | | | | | |
| C | CYS | | | | | | | | | | |
| V | VAL | 91 | 87 | 91 | 89.2 | 181 | 178.36 | 101 | 100.98 | 201 | 199.53 |
| M | MET | 1 | 1.18 | 1 | 1 | 1 | 1.05 | 1 | 1 | 1 | 1.15 |
| W | TRP | | | | | | | | | | |
| F | PHE | | | | | | | | | | |
| I | ILE | | | | | | | | | | |
| L | LEU | 2 | 2.01 | 2 | 2.09 | 2 | 2.75 | 2 | 2.26 | 2 | 2.12 |
| K | LYS | | | 10 | 9.67 | 10 | 9.99 | | | | |
| P | PRO | 51 | 51.31 | 51 | 51.2 | 101 | 106.23 | 51 | 51.09 | 101 | 100.05 |

**Table S3.** $pK_a$ values of glutamate residues in $V_2EV_2$ and V4K1- $V_2EV_2$ as a function of temperature, determined by acid-base titration. Reported values correspond to the mean ± standard deviation (*N* = *3*).

| | Temperature | | | | | | | | |
|---|---|---|---|---|---|---|---|---|---|
| | 5°C | 10°C | 15°C | 25°C | 40°C | 45°C | 50°C | 55°C | 60°C |
| $V_2EV_2$ | 4.24 ± 0.02 | 4.29 ± 0.05 | 4.30 ± 0.02 | 4.33 ± 0.03 | 4.41 ± 0.10 | 4.51 ± 0.12 | 4.55± 0.14 | - | 4.70 ± 0.14 |
| $V_2KV_2$-$V_2EV_2$ | 4.23 ± 0.15 | 4.24 ± 0.21 | 4.29 ± 0.08 | 4.43 ± 0.09 | 4.50 ± 0.15 | 4.62 ± 0.02 | 4.45 ± 0.07 | 4.31 ± 0.19 | 4.15 ± 0.14 |

**Table S4.** Enthalpy (ΔH°) and entropy (ΔS°) values obtained by Van't Hoff analysis.

| IDPP | ΔH° ($kcal\ mol^{-1}$) (T < $T_t$) | (T > $T_t$) | ΔS° ($cal\ mol^{-1}\ K^{-1}$) (T < $T_t$) | (T > $T_t$) |
|---|---|---|---|---|
| $V_2EV_2$ | -1.61 ± 0.35 | -6.69 ±0. 68 | -25.39 ± 1.23 | -41.56 ± 2.11 |
| $V_2KV_2$-$V_2EV_2$ | -3.44 ± 0.44 | 14.85 ± 0.26 | -33.5 ± 1.5 | 9.43 ± 0.80 |

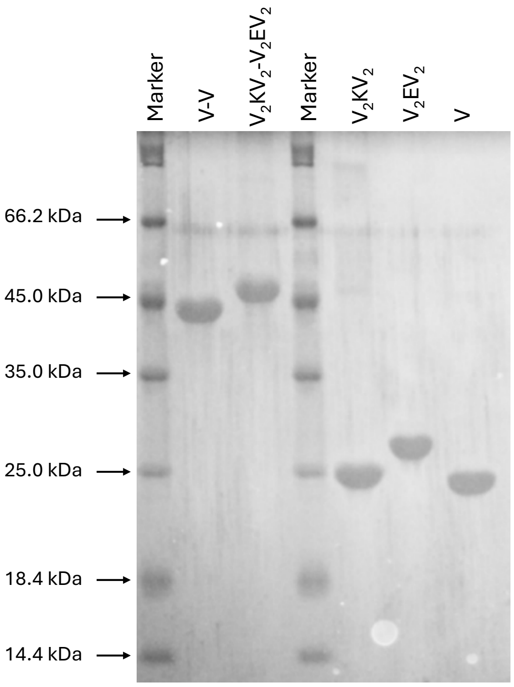


**Figure S1. SDS-PAGE electrophoresis of the recombinant IDPP library**.

Lanes 1 and 4 correspond to Pierce™ Unstained Protein MW Marker (ThermoFisher Scientific), with molecular weights indicated on the left. Lanes 2, 3, 5, and 6 show the purified IDPPs. The gel was stained with $CuCl_2$. The presence of single, sharp bands confirms the high purity and homogeneity of the expressed proteins.

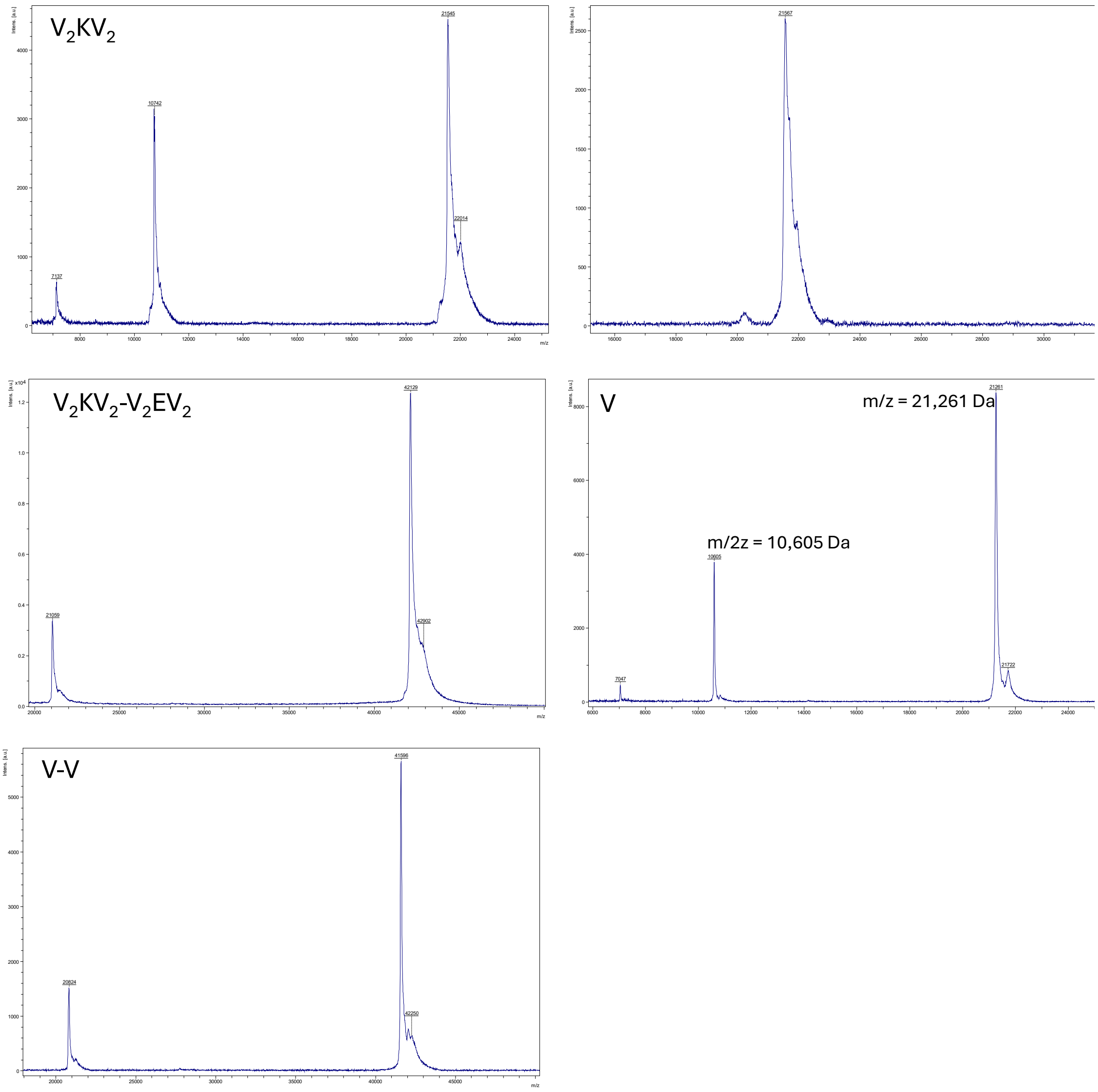


**Figure S2. MALDI-TOF mass spectra of the IDPPs.**

Spectra correspond to (a) $V_2KV_2$, (b) $V_2EV_2$, (c) $V_2KV_2$- $V_2EV_2$ diblock, (d) V [(VPGVG)50], and (e) V-V diblock [(VPGVG)100]. Minor peaks at approximately half the molecular weight correspond to doubly charged species (m/2z).

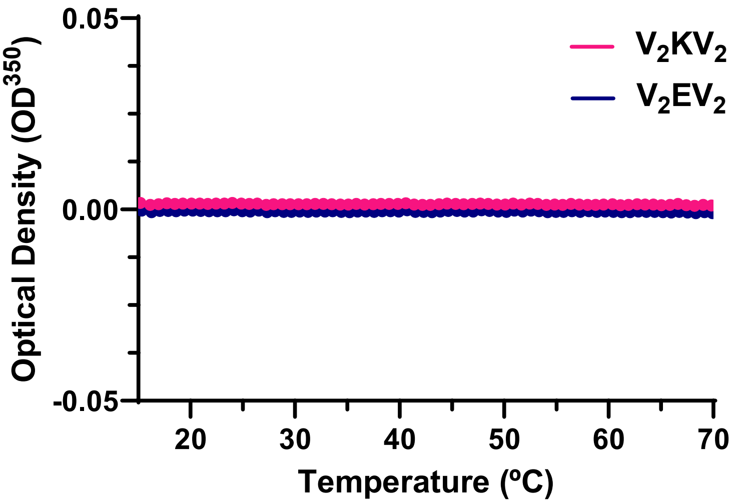


**Figure S3. Turbidity profiles of individual $V_2EV_2$ and $V_2KV_2$ IDPPs at pH 7.**

UV/Vis turbidity measurements as a function of temperature at 50 µM in ultrapure water (for $V_2EV_2$ and $V_2KV_2$ alone). No detectable LCST transition was observed under the tested conditions, indicating the absence of phase separation for either charged IDPP individually in water.

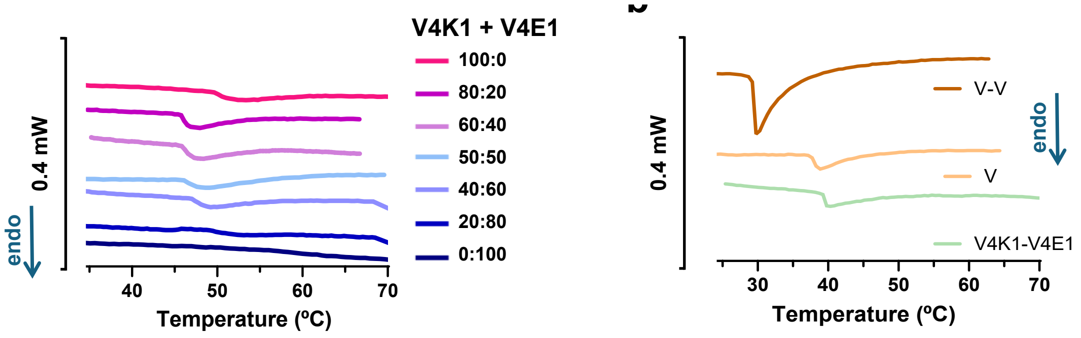


**Figure S4. Representative DSC thermograms measured at 5% (w/v) in PBS.** (a) Thermograms of mixtures of $V_2KV_2$ and $V_2EV_2$ at different molar ratios (from 0:100 to 100:0). (b) Thermograms of the diblock copolymer $V_2KV_2$-$V_2EV_2$ and the non-charged IDPPs V $(VPGVG)_{50}$ and V–V $(VPGVG)_{100}$.

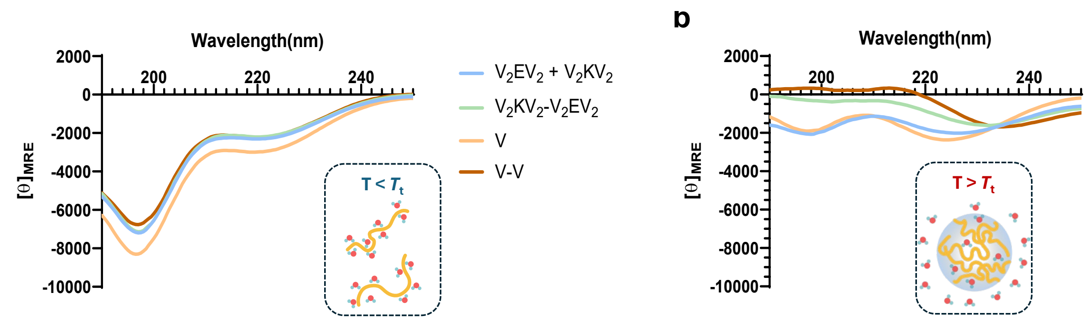


**Figure S5. Secondary structure analysis of the IDPPs.**

Below the transition temperature (T < $T_t$) (a), all IDPPs exhibit disordered conformations characterized by a strong negative minimum near 195 nm and a weak shoulder around 220 nm. Above the $T_t$ (b), the spectra change markedly: the minimum at 195 nm decreases, indicating reduced random-coil content, while the shoulder near 220 nm becomes more intense and slightly shifts toward 230 nm depending the composition.

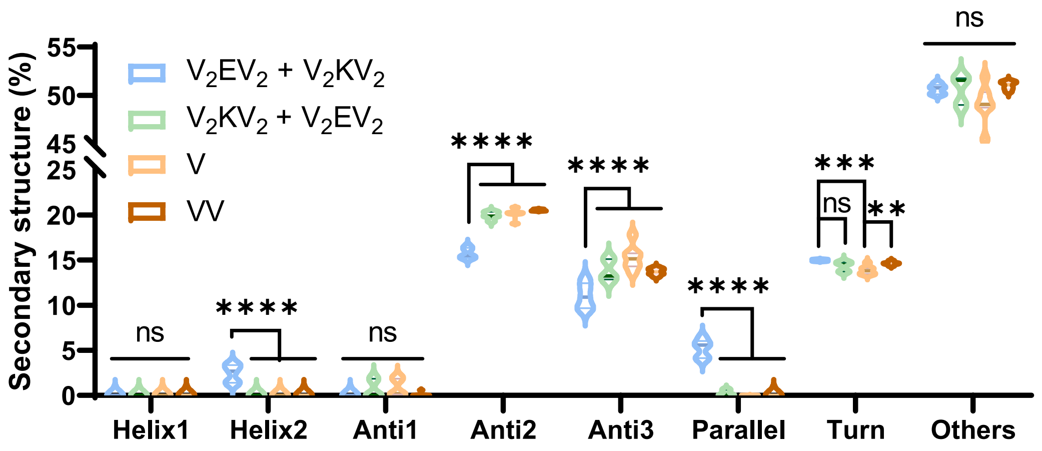


**Figure S6. BeStSel deconvolution of the secondary structures found in the IDPP condensates (above the transition temperature).**

Deconvolution of CD spectra using the BeStSel algorithm revealed that all condensates display mixed conformational content rather than a purely disordered signature. The spectra consistently show ~49–51% "Others" (random coil) and ~13.9–15.0% turns, together with a substantial β-structure contribution of ~32–36%. The β fraction is dominated by antiparallel components (Anti2 + Anti3), increasing from ~26.8% in the physically mixed $V_2KV_2$ + $V_2EV_2$ system to ~34.5% in the covalent $V_2KV_2$-$V_2EV_2$ diblock, ~36.3% in V, and ~34.3% in V-V. Notably, only the $V_2KV_2$ + $V_2EV_2$ mixture exhibits a parallel β component (~5.2%) and a minor helical contribution (~2.5%), both absent in $V_2KV_2$-$V_2EV_2$, and the hydrophobic analogues, V, and V-V (≤0.3% parallel; 0% helix). These results indicate that although the condensates remain predominantly disordered, they contain significant β-structures whose orientation depends on polymer architecture: physical mixing ($V_2EV_2$ + $V_2KV_2$) promotes a modest parallel arrangement and trace helicity, whereas covalent linkage ($V_2KV_2$-$V_2EV_2$) or longer chain length (V-V) favor antiparallel β enrichment.

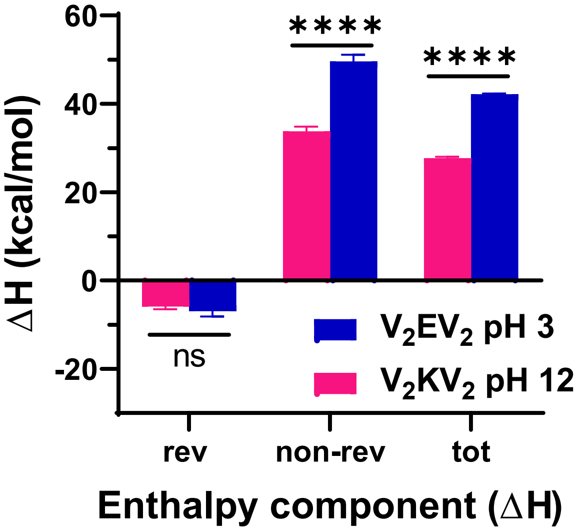


**Figure S7. Temperature-modulated DSC (TM-DSC) deconvolution of enthalpic components for $V_2EV_2$ and $V_2KV_2$ (5% w/v in water).** Samples were adjusted to pH values below or above the side-chain $pK_a$ to suppress electrostatic charges ($V_2EV_2$, pH 3; $V_2KV_2$, pH 12). Non-reversing (slow) and reversing (fast) enthalpic contributions are shown, highlighting the effect of protonation state on cooperative dehydration during LLPS. (*N=3*)

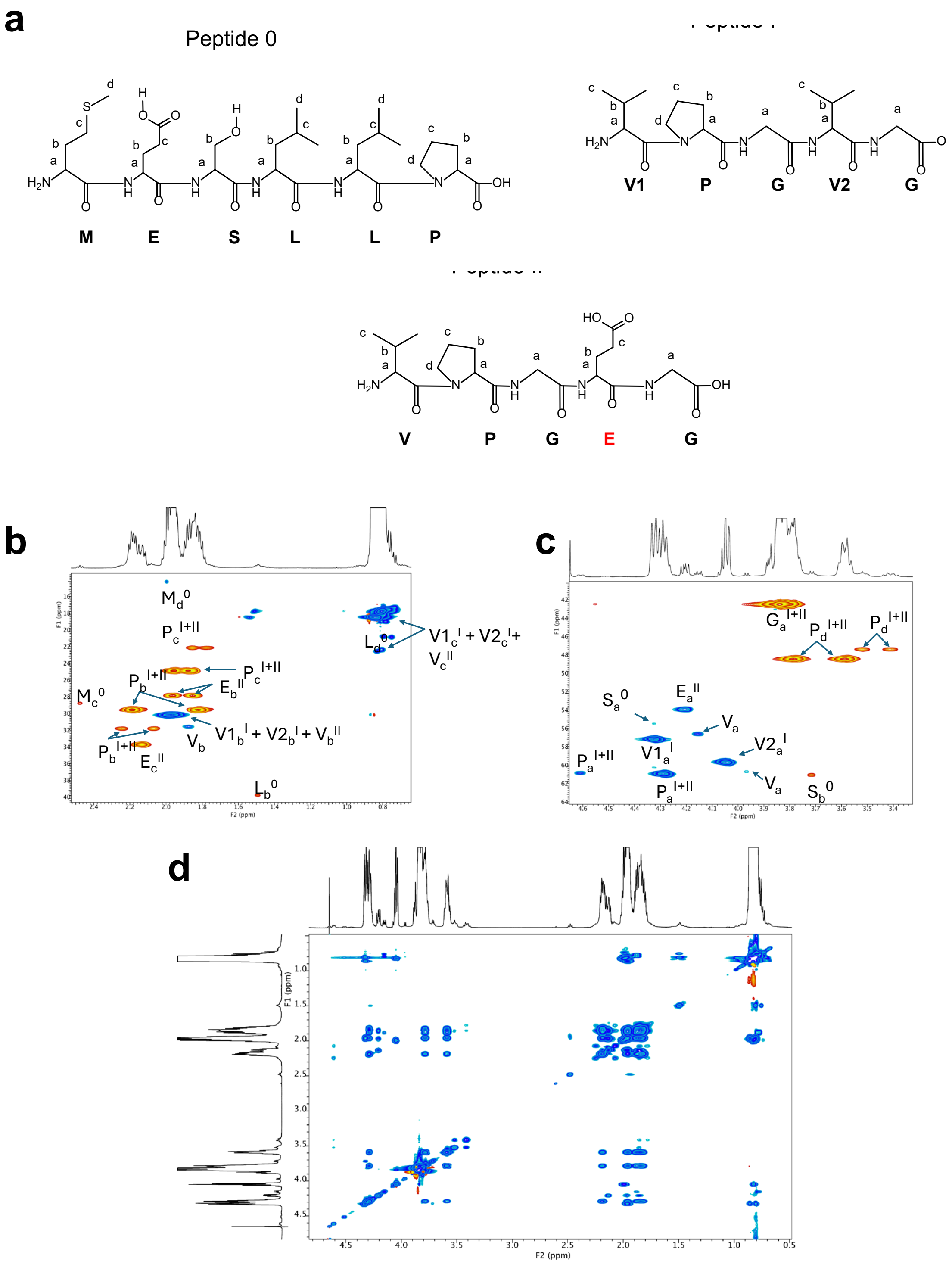


**Figure S8. NMR characterization of the $V_2EV_2$ system in solution.**

(a) Proton-labeling scheme used for residue assignments. Each proton is annotated as $X_y^{(n)}$, where X denotes the amino acid residue, *y* denotes the proton position within the residue ((a–e, corresponding to Cα–Cε)), and *(n)* indicates the peptide monomer (0: MESLLP leader; I: VPGVG; II: VPGEG). Because the spectra were acquired in $D_2O$, all exchangeable N–H protons, including backbone amides, undergo rapid H/D exchange and are therefore not detected.

(b) $^{1}H$–$^{13}C$ HSQC spectrum (0.6-2.6 ppm, $^{1}H$ dimension). This region contains $-CH_3$ groups from Val and Leu ($V_c^{I+II}$, $L_d^0$), methylene ($-CH_2-$)groups (e.g., $V_b^{I+II}$, $E_{b,c}^{II}$, $M_{c,d}^{0}$, $P_{b,c}^{I+II}$).

(c) $^{1}$H–$^{13}$C HSQC spectrum (3.0-4.8 ppm) displaying the region that is dominated by Cα–H correlations from Val, Pro, Glu, Lys, and Ser ($V_a^{I,II}$, $P_a^{I+II}$, $E_a^{II}$, $S_a^{0}$). This region also includes the intense Cα–CH₂ signals of Gly ($G_a^{I+II}$) and those corresponding to Ser Cβ ($S_b^{0}$) and Pro Cδ ($P_d^{I+II}$).

(d) $^{1}$H–$^{1}$H zTOCSY spectrum below the $T_t$ revealing intraresidue spin-system correlations in agreement with HSQC assignments.

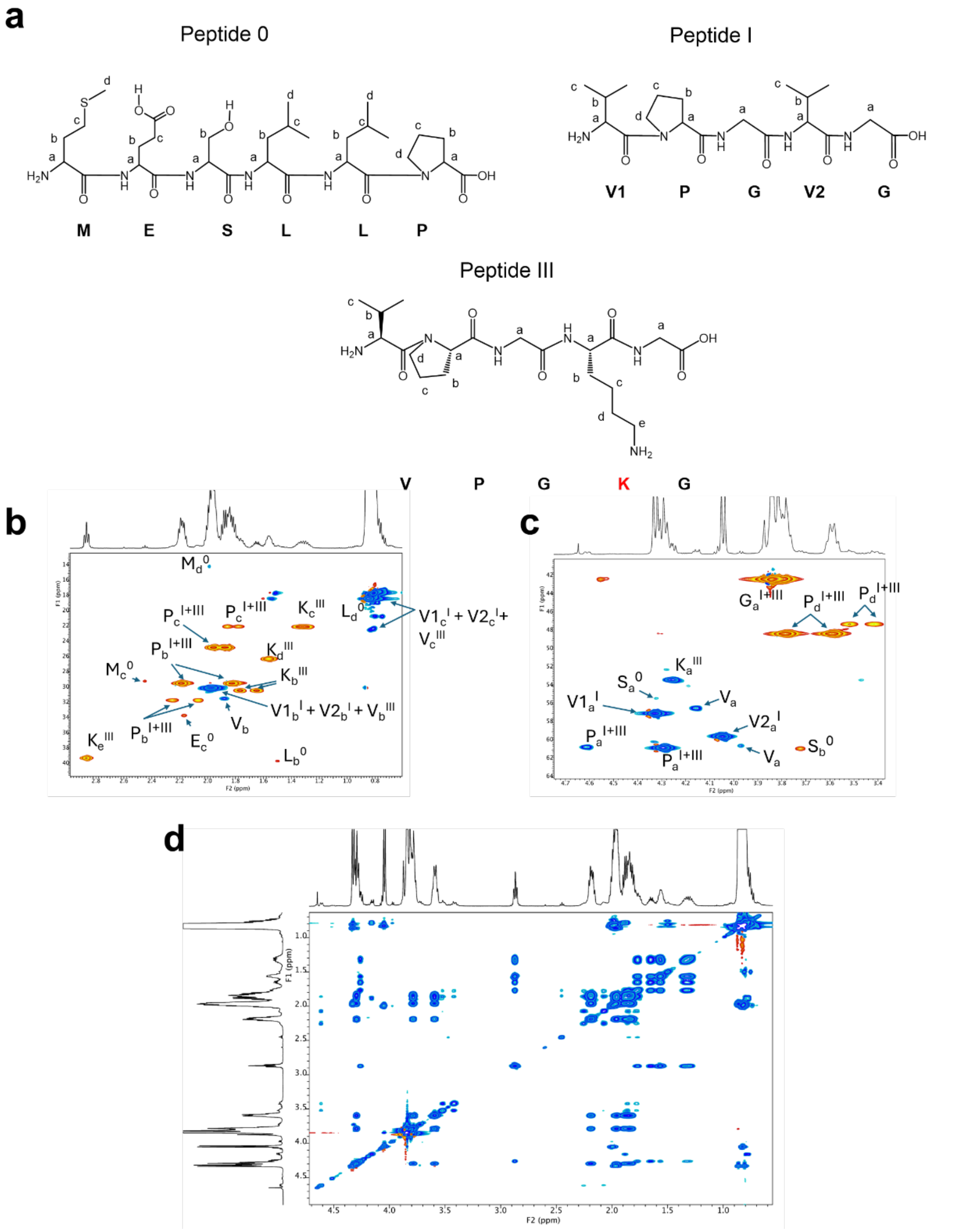

**Figure S9. NMR characterization of the $V_2KV_2$ in solution.**

(a) Proton-labeling scheme used for residue assignments. Each proton is annotated as $X_y^{(n)}$, where X denotes the amino acid residue, *y* denotes the proton position within the residue (a–e), and *(n)* indicates the peptide monomer (0: MESLLP leader; I: VPGVG; III: VPGKG).

(b) $^1H$–$^{13}C$ HSQC spectrum (0.6-2.6 ppm, $^1H$ dimension). This region contains -$CH_3$ groups from Val and Leu ($V_c^{I+III}$, $L_d^0$), the characteristic Lys side-chain methylenes ($K_{b,c,d,e}^{III}$), as well as -$CH_2$- signals from Met and Pro ($M_{c,d}^0$, $P_{b,c}^{I,III}$).methylene (-$CH_2$-)groups (e.g., $V_b^{I+III}$, $M_{c,d}^0$, $P_{b,c}^{I+III}$).

(c) $^1H$–$^{13}C$ HSQC spectrum (3.0-4.8 ppm) displaying the region that is dominated by Cα–H correlations from Val, Pro, Glu, Lys, and Ser ($V_a^{I,III}$, $P_a^{I+III}$, $S_a^0$). This region also includes the intense Cα–$CH_2$ signals of Gly ($G_a^{I+III}$) and those corresponding to Ser Cβ ($S_b^0$) and Pro Cδ ($P_d^{I+III}$).

(d) $^1H$–$^1H$ zTOCSY spectrum below the $T_t$ revealing intraresidue spin-system correlations in agreement with HSQC assignments.

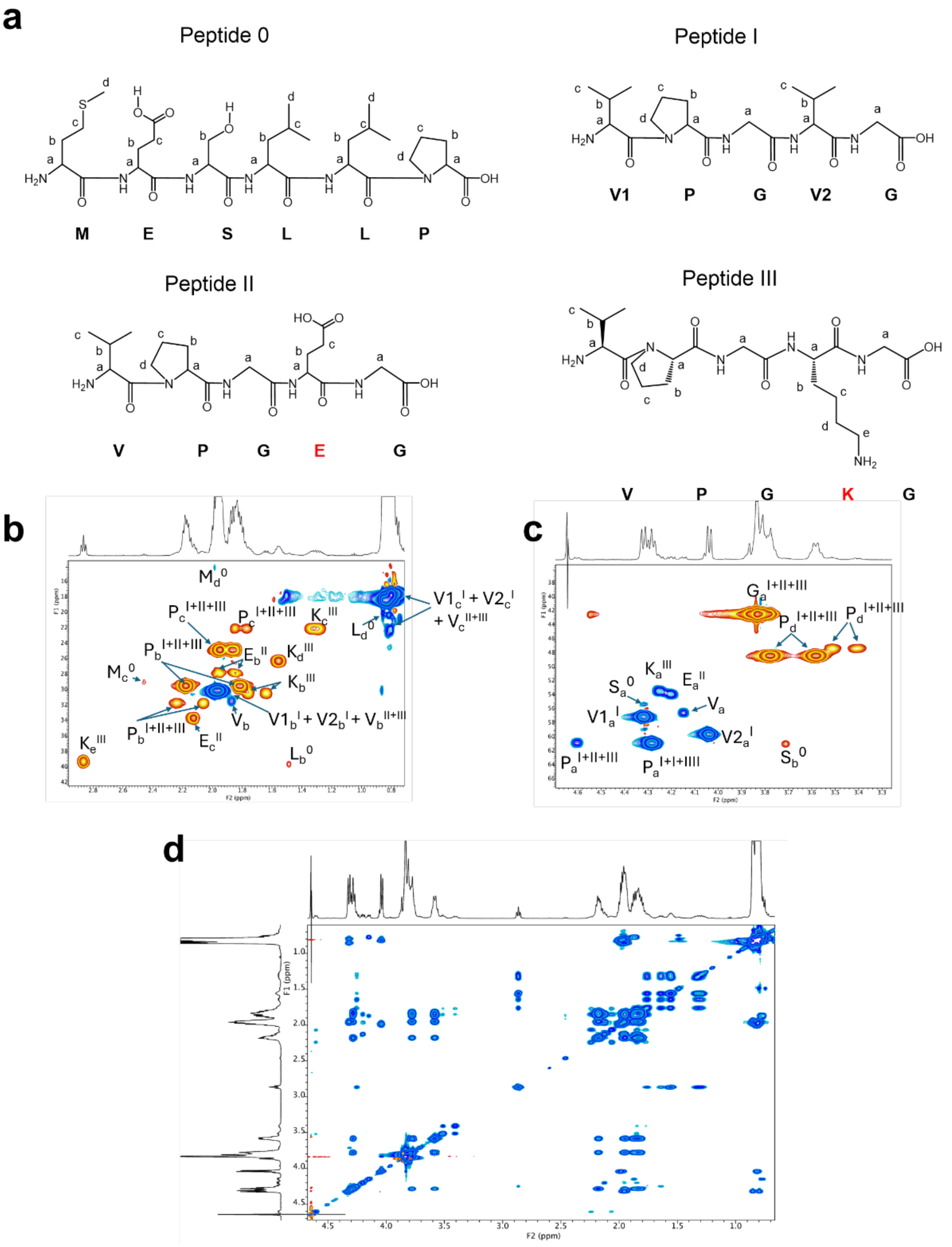


**Figure S10. NMR characterization of the $V_2KV_2$–$V_2EV_2$ system in the soluble state (below the $T_t$, T = 25 °C).**

(a) Proton-labeling scheme used throughout the NMR analysis. Each signal is annotated as $X_y^{(n)}$, where X denotes the amino acid residue, *y* denotes the proton position within the residue (a–e), and *(n)* indicates the peptide monomer (0: MESLLP leader; I: VPGVG; II: VPGEG; III: VPGKG). Only protons directly bound to carbon atoms are labeled.

(b) $^1$H–$^{13}$C HSQC spectrum (0.6-3.0 ppm, $^1$H dimension) showing -$CH_3$ groups from Val and Leu ($V_c^{I+II+III}$, $L_d^0$), methylene (-$CH_2$-)groups from Lys, Glu, Met, and Pro (e.g., $K_{b-e}^{III}$, $E_{b,c}^{II}$, $M_{c,d}^0$, $P_{b,c}^{I+II+III}$), and additional Cβ resonances from the peptide 0-I.

(c) $^1$H–$^{13}$C HSQC spectrum (3.0-4.8 ppm) displaying the region that is dominated by Cα–H correlations from Val, Pro, Glu, Lys, and Ser ($V_a^{I,II,III}$, $P_a^{I+II+III}$, $E_a^{II}$, $K_a^{III}$, $S_a^0$). This region also includes the intense Cα–CH₂ signals of Gly ($G_a^{I+II+III}$) and those corresponding to Ser Cβ ($S_b^0$) and Pro Cδ ($P_d^{I+II+III}$).

(d) $^1$H–$^1$H zTOCSY spectrum below the $T_t$ revealing intraresidue spin-system correlations in agreement with HSQC assignments.

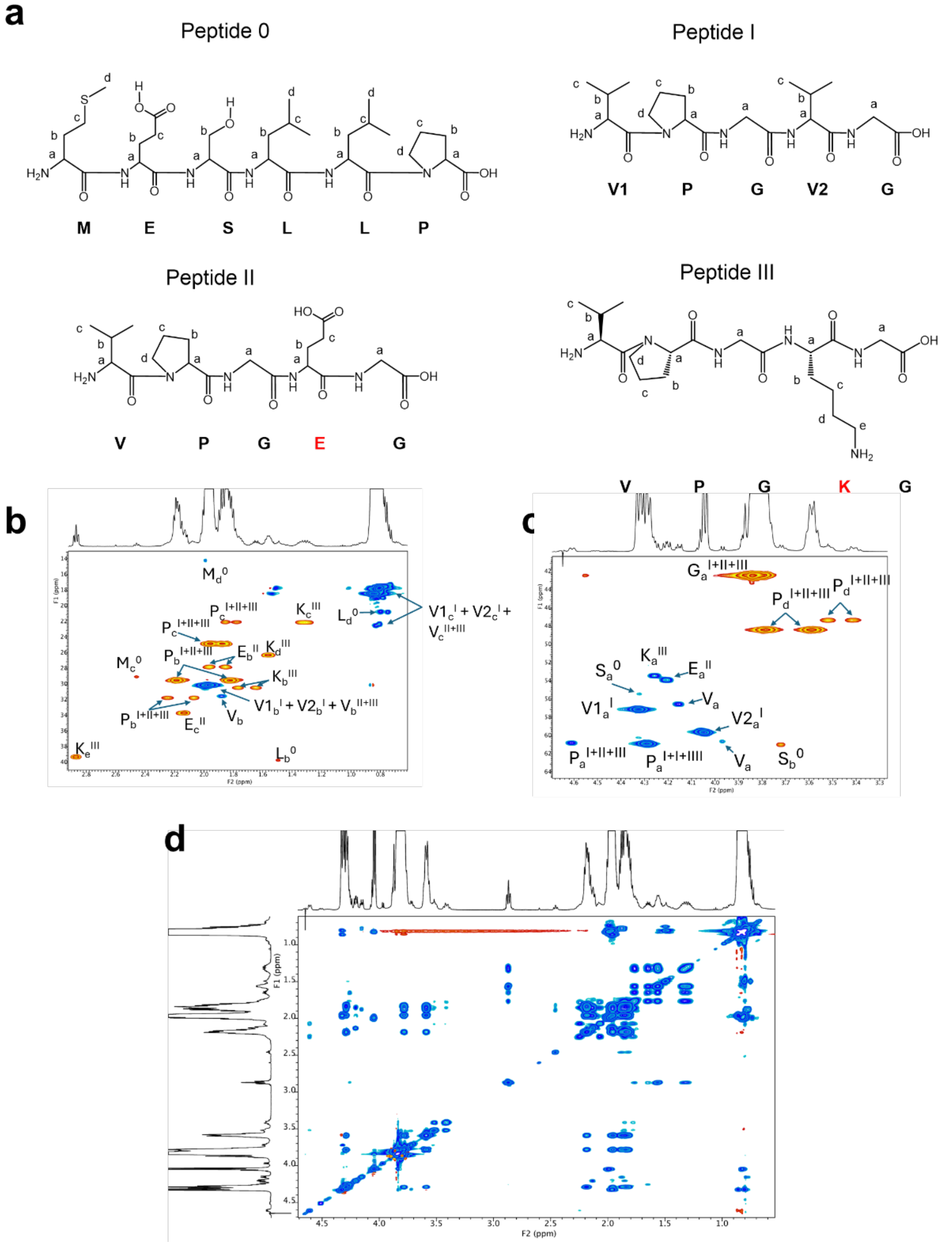

**Figure S11. NMR characterization of the $V_2EV_2$ + $V_2KV_2$ system in the soluble state (below the $T_t$, T = 25 °C).**

(a) Proton-labeling scheme used throughout the NMR analysis. Each signal is annotated as $X_y^{(n)}$, where X denotes the amino acid residue, *y* denotes the proton position within the residue (a–e), and *(n)* indicates the peptide monomer (0: MESLLP leader; I: VPGVG; II: VPGEG; III: VPGKG). Only carbon-bound protons are labeled. Because measurements were performed in $D_2O$, exchangeable protons (amide NH and the Lys $NH_3^+$ protons) are absent due to rapid H/D exchange.

(b) $^1$H–$^{13}$C HSQC spectrum (0.6-3.0 ppm, $^1$H dimension) showing -$CH_3$ groups from Val and Leu ($V_c^{I+II+III}$, $L_d^0$), methylene (-$CH_2$-)groups from Lys, Glu, Met, and Pro (e.g., $K_{b-e}^{III}$, $E_{b,c}^{II}$, $M_{c,d}^0$, $P_{b,c}^{I+II+III}$), and additional Cβ resonances from the peptide 0-I.

(c) $^1$H–$^{13}$C HSQC spectrum (3.0-4.8 ppm) displaying the region that is dominated by Cα–H correlations from Val, Pro, Glu, Lys, and Ser ($V_a^{I,II,III}$, $P_a^{I+II+III}$, $E_a^{II}$, $K_a^{III}$, $S_a^0$). This region also includes the intense Cα–$CH_2$ signals of Gly ($G_a^{I+II+III}$) and those corresponding to Ser Cβ ($S_b^0$) and Pro Cδ ($P_d^{I+II+III}$).

(d) $^1$H–$^1$H zTOCSY spectrum below the $T_t$ revealing intraresidue spin-system correlations in agreement with HSQC assignments.

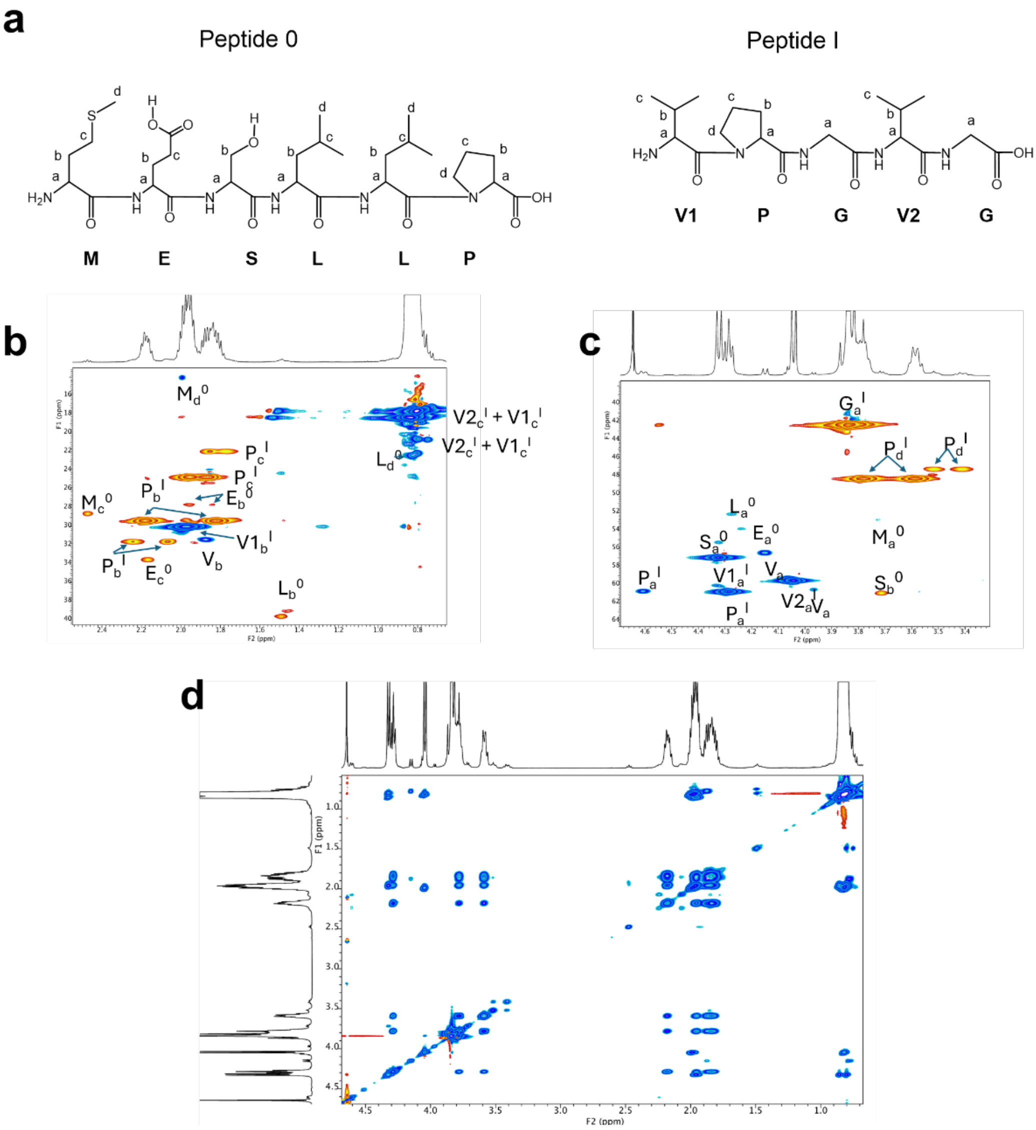


**Figure S12. NMR characterization of the V polymer in the soluble state (below the $T_t$, T = 25 °C).**

(a) Proton-labeling scheme used throughout the NMR analysis. Each signal is annotated as $X_y^{(n)}$, where X denotes the amino acid residue, *y* denotes the proton position within the residue (a–e), and *(n)* indicates the peptide monomer (0: MESLLP leader; I: VPGVG). Only protons directly bound to carbon atoms are labeled.

(b) $^1H$–$^{13}C$ HSQC spectrum (0.6-2.6 ppm, $^1H$ dimension) showing -$CH_3$ groups from Val and Leu ($V_c^I$, $L_d^0$), the β-methylene (Cβ) of Val ($V_b$), and the Cγ and Cδ methylenes of Pro ($P_{b,c}$), in addition to the methylene (-$CH_2$-)groups from the leader peptide 0.

(c) $^1H$–$^{13}C$ HSQC spectrum (3.0-4.8 ppm) displaying the region that is dominated by Cα–H correlations from Val, Pro, Glu, Lys, and Ser ($V_a^I$, $P_a^I$, $S_a^0$). This region also includes the intense Cα–$CH_2$ signals of Gly ($G_a^I$) and the only two -$CH_2$- resonances appearing above 3 ppm, corresponding to Ser Cβ ($SS_b^0$) and Pro Cδ ($P_d^I$).

(d) $^1H$–$^1H$ zTOCSY spectrum below the $T_t$ revealing intraresidue spin-system correlations in agreement with HSQC assignments.

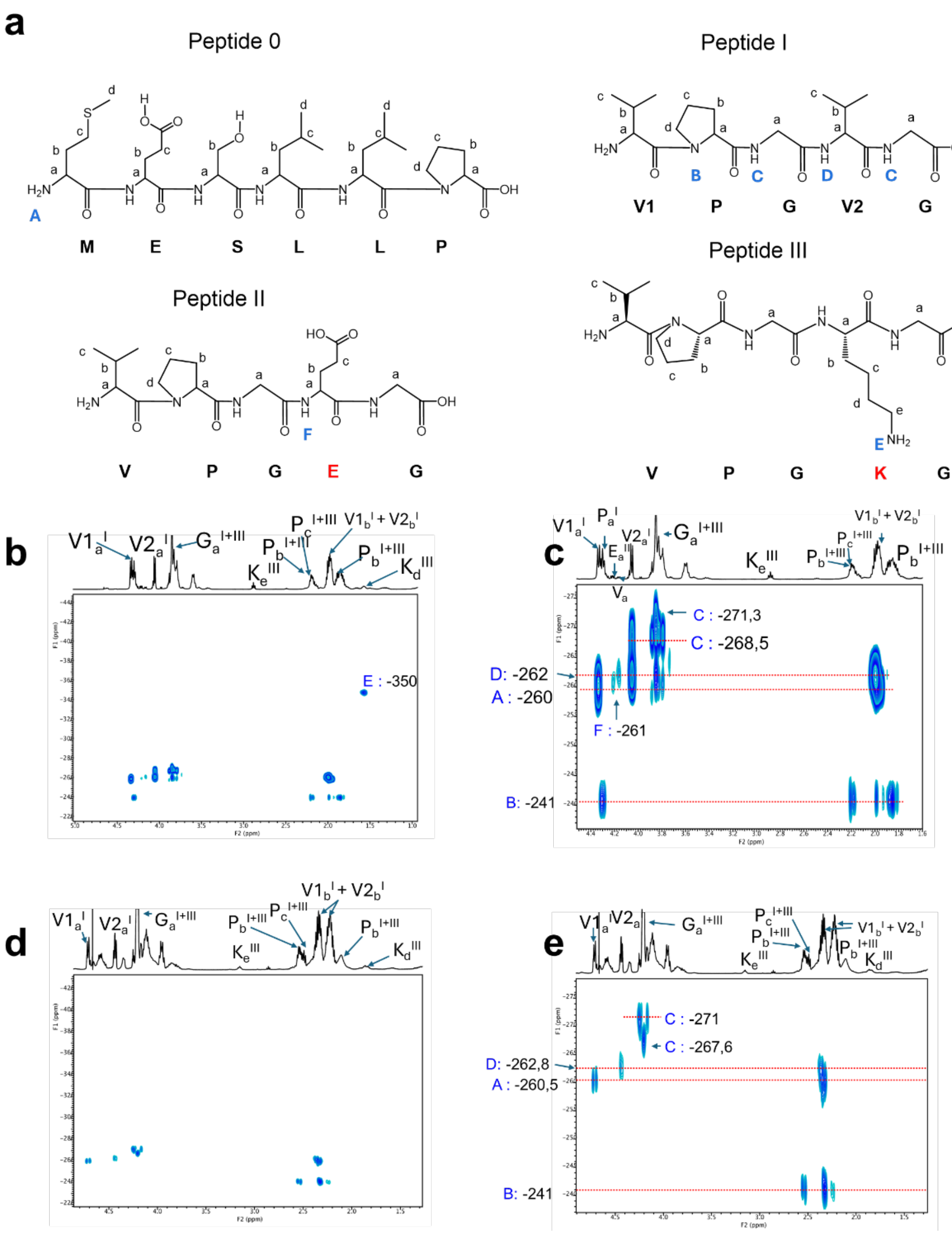

**Figure S13. $^{1}H$–$^{15}N$ HMBC characterization of the $V_2E\ V_2 + V_2K\ V_2$ system.**

(a) Proton-nitrogen labeling scheme used for $^{15}N$ assignments. Each nitrogen correlation is annotated using blue uppercase letters (A–F), corresponding to distinct $^{15}N$ chemical environments arising from N-terminus, Lys primary amine, and backbone amide positions within the two-component ($V_2E\ V_2 + V_2K\ V_2$) system.

(b) 2D $^{1}H$-$^{15}N$ HMBC spectra acquired below the $T_t$ (25 °C), where the mixture remains fully solvated.

(c) Magnified view (zoom-in) of panel (b), highlighting the nitrogen region between –230 and –280 ppm. This zoom resolves overlapping correlations and improves visualization of nitrogen signals A, B, C, D and F.

(d) $^{1}H$–$^{15}N$ HMBC spectrum above the $T_t$ (60 °C), where $V_2E\ V_2 + V_2K\ V_2$ system is condensed. (e) Zoom-in of panel (d), resolving the remaining nitrogen signals in the –230 to –280 ppm region.

Only nitrogen A–D remained observable. Cross-peaks corresponding to Glu amide nitrogen (F) and the Lys ε-amine (E) were no longer detected.

The loss of signals E and F is attributed both to natural-abundance limitations of $^{15}N$, together with severe line broadening due to rapid transverse relaxation (short $T_2$) associated with the restricted molecular motion and increased microviscosity in the dense phase. These effects suppress long-range $^{1}H$-$^{15}N$ correlations for dynamically sensitive nitrogen environments.

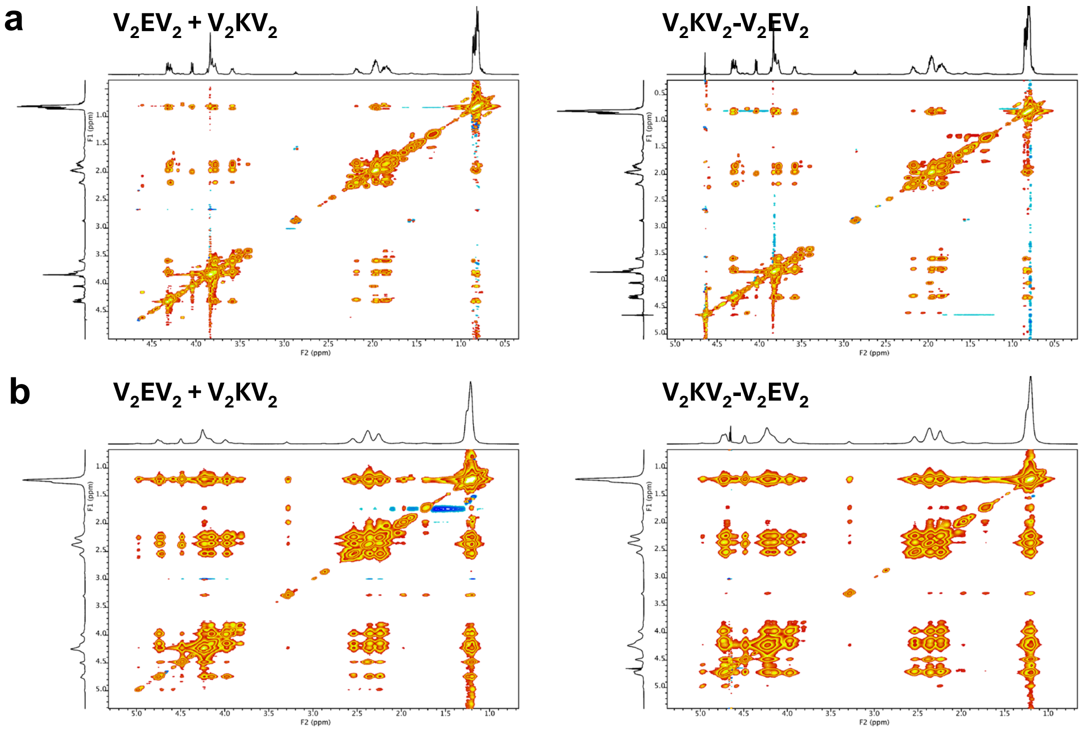


**Figure S14. 2D $^{1}H$-$^{1}H$ NOESY spectra of charged IDPPs measured below (a) and above (b) the transition temperature ($T_t$).**

2D NOESY spectra of the physical mixture $V_2EV_2$ + $V_2KV_2$ and the covalently bonded $V_2KV_2$-$V_2EV_2$ acquired at 25 °C (T < $T_t$) and 60 °C (T > $T_t$) in $D_2O$ to monitor temperature-induced changes in proton-proton correlations. Despite the clear temperature-dependent changes observed in the NOESY spectra, no unambiguous cross-peak indicative of Glu-Lys side-chain association could be identified in the condensed phase. This negative result likely arises from two factors. First, the β-methylene protons of Glu and the ε-methylene protons of Lys might be separated by distances that exceed the ~5 Å spatial limit required for observable NOE build-up, making direct through-space detection intrinsically unlikely even if salt-bridge formation occurs. Second, above the $T_t$ the spectra undergo severe line broadening due to reduced molecular mobility in the condensed phase, which further suppresses NOE intensities and leads to overlap with nearby aliphatic resonances.

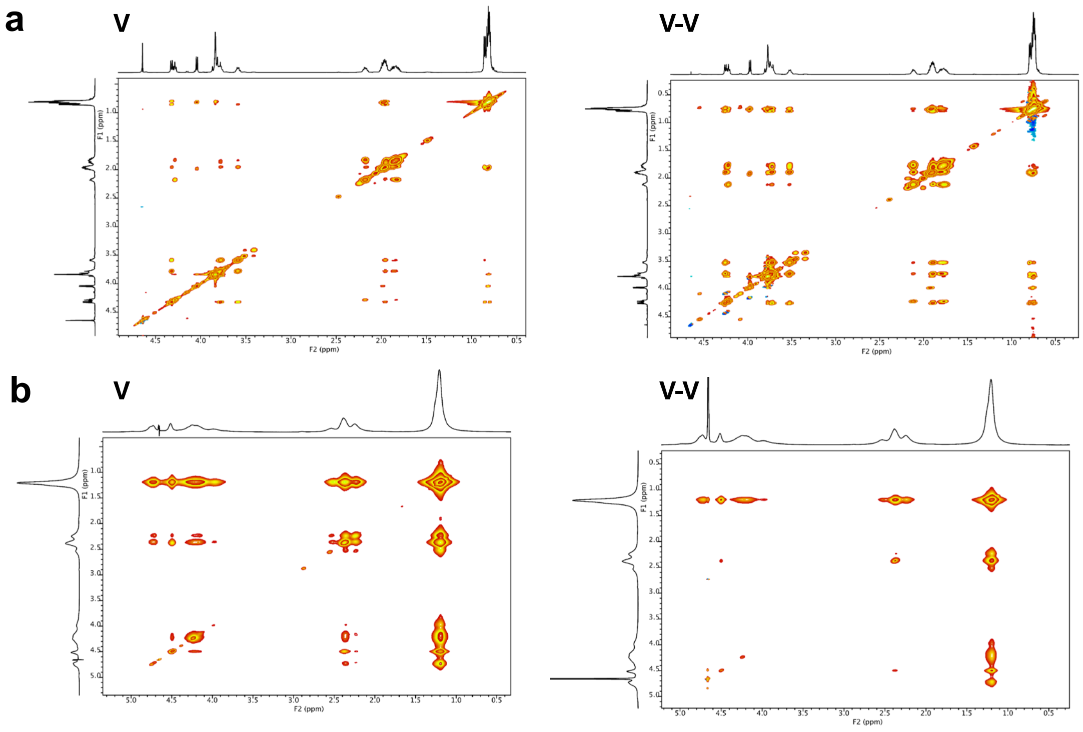


**Figure S15. 2D $^1$H-$^1$H NOESY spectra of hydrophobic IDPPs measured below (a) and above (b) the transition temperature ($T_t$).**

Spectra of V and V-V IDPPs acquired at 25 °C (T < $T_t$) and 60 °C (T > $T_t$) under identical conditions to the charged systems. Below the $T_t$, both samples display sharp, well-resolved cross-peaks characteristic of flexible, hydrated chains in the soluble state. Above the $T_t$, the spectra exhibit a pronounced broadening of cross-peaks, more evident for the V-V, consistent with reduced molecular mobility within the condensate. This broadening reflects increased chain entanglement and slower tumbling associated with phase condensation, rather than specific side-chain interactions.

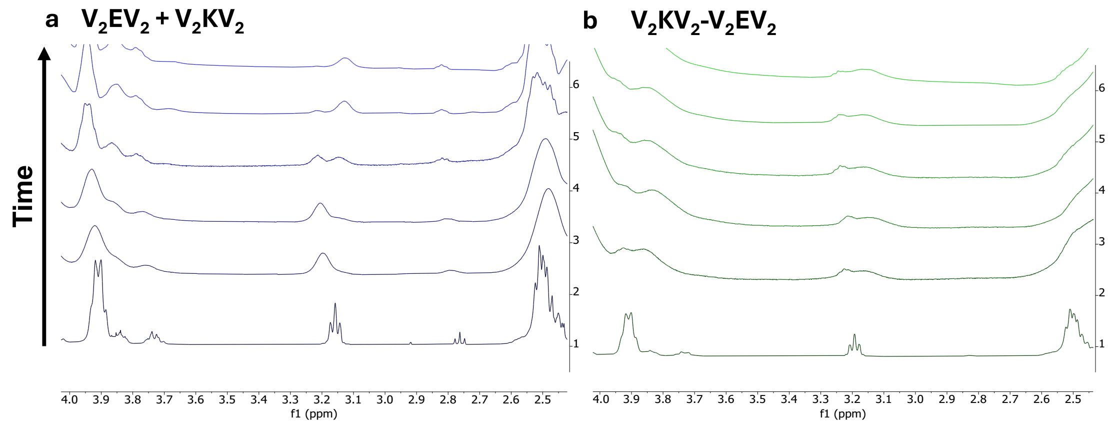


**Figure S16. Representative $^1$H-NMR spectra of charged IDPP condensates (expanded region, 2.5–4.0 ppm) recorded at different times after surpassing the transition temperature (T > $T_t$).**

Spectra of (a) the physical mixture $V_2EV_2$ + $V_2KV_2$ and (b) the covalently bonded $V_2KV_2$-$V_2EV_2$ were collected at 60 °C at selected time points (0, 1, 2, 8, 43, and 200 min). The expanded region focuses on the ε-methylene protons of lysine, which serve as sensitive reporters of local microenvironment and ionization state. Over time, a progressive broadening and attenuation of the signals is observed, consistent with aggregation and reduced molecular mobility within the condensed phase. Around 3.2-3.1 ppm, two partially resolved populations appear: one corresponding to free Lys–$NH_3^+$ groups and the other to Lys-$NH_2$-H-Glu salt-bridge formation, revealing the gradual formation of electrostatic interactions during condensate maturation.
